# Perceiving Instability: How Expectations Bias Sensorimotor Processing in Balance Control

**DOI:** 10.1101/2025.11.18.688816

**Authors:** Johnny V. V. Parr, Richard Mills, Elmar Kal, Adolfo M. Bronstein, Toby J. Ellmers

## Abstract

Maintaining balance requires rapid integration of sensory input with top-down sensorimotor predictions. Predictive coding frameworks propose that mismatches between expected and actual sensory input (“prediction errors”) drive perceptual inference. Although these frameworks have been highly influential in sensory neuroscience, it remains unclear whether they operate similarly in fast-acting sensorimotor systems such as human balance. Using electroencephalography (EEG) and discrete postural perturbations via support surface translations, we independently manipulated expectations and sensory input. Participants were primed to expect small or large perturbations, with occasional violations in which the delivered perturbation differed from expectation. We found that subjective perception of instability was shaped by expectation alone, regardless of perturbation magnitude. Similarly, pre-perturbation beta-band suppression and post-perturbation gamma activity tracked expected perturbation magnitude, with the latter strongly associated with expectation-induced perceptual biases (*r* = 0.69, *p_BON_* = 0.001). In contrast, both the balance N1 (a well-established stimulus-evoked cortical potential linked to the postural response) and *objective* behavioural markers of postural instability were determined primarily by perturbation magnitude. Crucially, neither perception nor early neural or behavioural markers appeared to encode prediction errors arising from expectation violations. Together, these findings identify boundary conditions on predictive coding in sensorimotor control, showing that during fast, reactive behaviour, perceptual inference may be shaped more strongly by expectations than by mismatch signals, even as early neural and motor responses remain driven primarily by sensory input.

**Significance Statement:** Perception is often explained through predictive coding, a framework in which the brain compares incoming sensory input with internal expectations and updates representations based on mismatches, or prediction errors. However, it remains unclear whether these perceptual inference processes operate similarly in fast, ecologically relevant sensorimotor systems, such as postural control. Our results show that top-down expectations shape perceptual experience and early cortical responses to postural perturbations, even when sensory input contradicts those expectations. Notably, these early neural markers showed limited sensitivity to prediction errors in this rapid, reactive context. Together, these findings identify conditions under which perceptual inference may rely more strongly on expectations than on sensory mismatch, refining predictive coding accounts of fast, reactive behaviour.

**Classification:** Biological Science (Neuroscience)

## Introduction

Recent advances in neuroscience have revealed that the brain operates as a ‘prediction machine’ (1). Rather than passively receiving sensory input from the periphery, the brain actively shapes perception by attempting to match incoming information to top-down expectations (2). This is useful, because it allows us to efficiently process and filter the vast amount of (often “noisy”) sensory information we encounter. But such processing shortcuts mean that perception can be inaccurately biased towards expectation, especially during scenarios that have thus far been highly predictable and/or when sensory input is ambiguous (with any resulting prediction errors thus explained away as “noise” rather than being used to update perceptual inference). Such expectation-based perceptual biases have been reported across sensory domains, including vision (3,4), audition (5,6), and pain (7–9).

It remains unknown, however, if the same expectation-based perceptual biases similarly affect sensory systems that are fast, automatic, and tightly coupled to motor output, such as balance control. Balance relies on rapid subcortical feedback loops (10,11) and continuous multisensory integration (12,13) to respond to postural disturbances. Although many of these disturbances (and associated postural adjustments) occur outside of conscious awareness (14,15), the balance system nonetheless gives rise to vivid subjective perceptions of (in)stability (16,17). But these perceptions can become biased and unreliable. Individuals frequently report feeling imbalanced even when postural stability nonetheless remains high (16,18) and, in some cases, these feelings persist despite clear sensory evidence to the contrary (19). This apparent dissociation between sensory evidence and subjective perception raises the possibility that prediction errors are weighted differently (or potentially suppressed entirely) during balance-related perceptual inference. This contrasts with classic frameworks that position prediction errors as the primary drivers of perceptual inference across both uni- and multi-sensory tasks (20). It is worth noting, however, that most of these frameworks are based on evidence from sensory tasks that do not inherently require rapid behavioural correction following a prediction error. This stands in contrast to balance control, in which an uncorrected prediction error could have catastrophic consequences (e.g., a fall). Determining whether predictive processing principles extend to balance perception is therefore critical for understanding how the brain negotiates the trade-off between perceptual accuracy and rapid motor action.

The present study examined how sensory signals interact with expectations (and violations of such, i.e. prediction errors) to shape perception and sensorimotor processing during a rapid, reactive balance task. Although prediction errors are known to guide longer-term balance behaviour (e.g., locomotor adaptation (21,22)), it remains unclear whether they shape within-trial balance perception or early sensorimotor processing that underlies immediate responses to postural instability crucial for preventing falls. Using electroencephalography (EEG), we measured neural responses during discrete postural perturbations whilst manipulating expectation about the size of the upcoming disturbance. Our results revealed that subjective perceptions of instability were strongly shaped by expectation: participants felt more unstable when anticipating large perturbations, regardless of actual perturbation received. We identified two temporally and spatially distinct neural markers that were modulated by expectation: anticipatory beta-band activity and post-perturbation gamma power (which was strongly and significantly associated with expectation-induced perceptual biases). In contrast, the balance N1, a well-established evoked early cortical response to instability (23,24), and early postural responses themselves were driven solely by perturbation size (i.e., sensory input) and unaffected by expectation. Crucially, none of these early signals showed strong sensitivity to prediction errors within the time window examined.

Our findings provide direct evidence that top-down expectations can bias how sensory signals are interpreted at both perceptual and neural levels during a rapid (and reactive) balance task. The absence of clear cortical markers tracking prediction error suggests that such signalling may not be uniformly expressed across all perceptual inference contexts, particularly during rapid, reactive motor control. Instead, while top-down expectations appear to dominate perceptual inference in fast-acting motor systems like balance, short-latency mechanisms (e.g., the N1) that track perturbation magnitude are preserved. Together, these parallel processes may help ensure that behaviour remains flexibly tuned to rapidly changing environments, minimising fall risk even when predictions are inaccurate and explicit prediction error processing may be too slow to support effective balance control.

## Results

Twenty-one neurotypical young adults experienced a series of anterior-posterior (AP) discrete balance perturbations. Perturbations were delivered under two blocks that manipulated expectation. In the Small-Expectation (SE) block of trials, participants were informed that they would encounter small perturbations (see Figure 1A) throughout the block of 60 trials. However, 15 of these perturbations violated expectations (i.e., they were ‘big’). These expectancy-violating perturbations were introduced on average every 2-3 trials and were never presented in succession. The Big-Expectation (BE) block of trials followed the same structure but with the perturbation sizes reversed. After certain expectation-confirming and -violating trials (see Methods section), participants also self-reported their levels of perceived instability and state anxiety during the preceding trial (17). Behavioural responses to each perturbation were operationalised as direction-specific (i.e., backwards) centre of pressure (COP) peak velocity.

**Figure 1.**
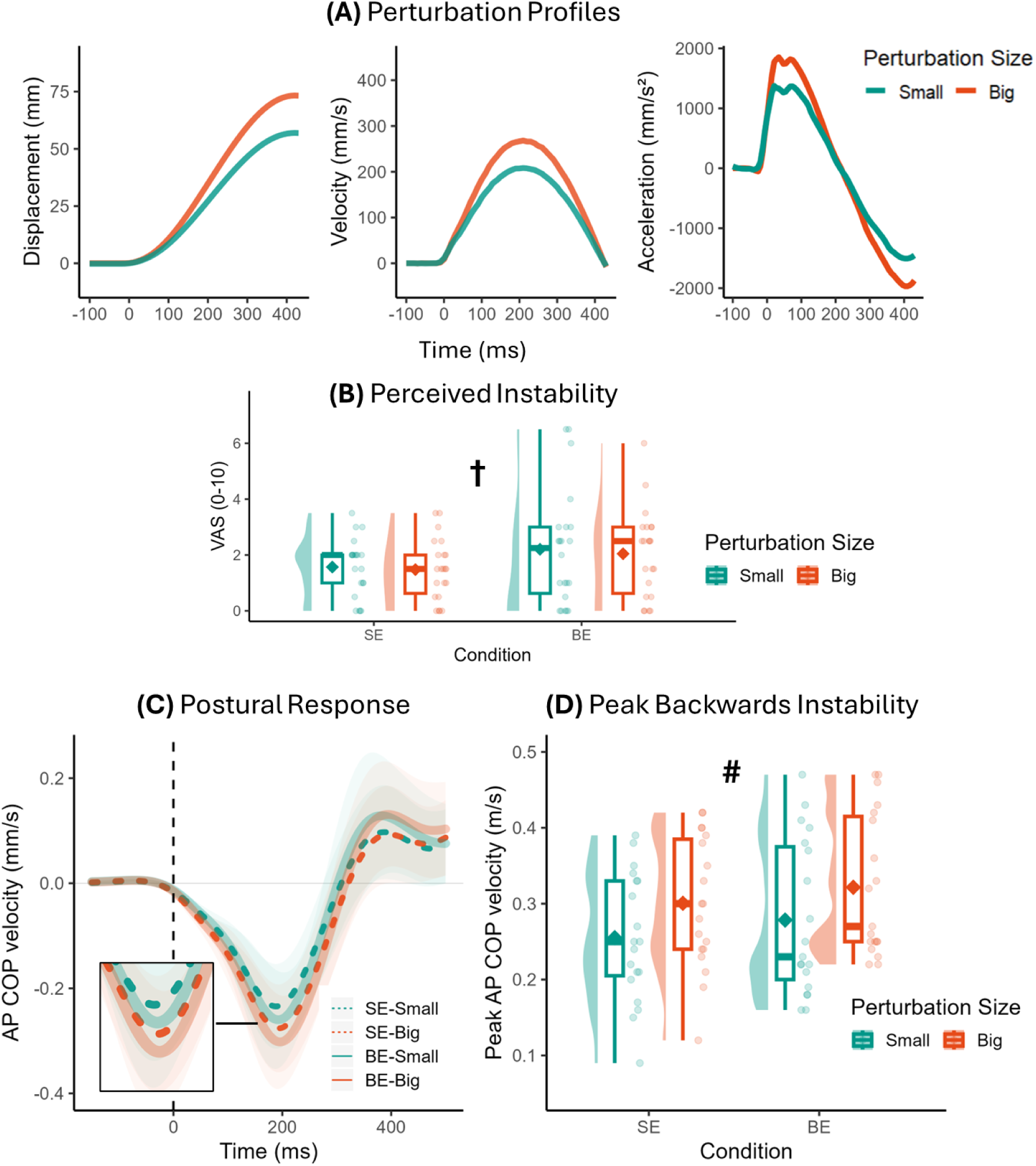
**A)** Displacement, velocity, and acceleration time-courses of the initial forward platform translation for each perturbation size. **B)** Subjective balance perception (perceived instability). **C)** Grand averages (±SD) of anterior-posterior (AP) centre of pressure (COP) velocity time-course following perturbation onset. **D)** Objective balance outcomes of peak instability (peak backwards centre of pressure COP velocity) across the four trial types. Note, SE reflects Small-Expectation trials whilst BE reflects Big-Expectation trials (e.g., ‘SE-Small’ indicates that participants expected a small perturbation and received this). Condition means for boxplots on 1B and 1D are denoted with ◊. Significant main effects of Expectation (†) and Perturbation Size (#) are denoted on the Figures where relevant. Please see the Supplementary Material 1 for extended data visualisation of within-subject changes from Figures 1B (Supplementary Figure S1.1) and 1D (Supplementary Figure S1.2) as a function of expectation and perturbation size.

We assessed EEG throughout the experiment. Specifically, we analysed pre-perturbation beta activity, given that lower pre-stimulus beta activity is reliably linked to enhanced sensory sensitivity (25,26) and might thus be sensitive to top-down expectation. We then also analysed perturbation-evoked responses: the balance N1 (a negative response occurring across the supplementary motor area ∼150ms after an external balance perturbation (27,28)) and post-perturbation gamma activity. Whilst balance N1 amplitude typically scales with perturbation magnitude (27,29), findings suggest that the N1 is attenuated when perturbations are temporally expected (23). Likewise, there is extensive research describing cortical gamma (>30 Hz) as a robust prediction error signal across tasks and sensory modalities (30,31). These outcomes will therefore provide insight into how prediction errors (e.g., trials that violate expectations) shape cortical processing of balance. Across outcomes, the presence of a significant Expectation×Perturbation Size interaction would indicate that the behavioural, perceptual or neural outcome significantly differed on trials that violated expectations (e.g., expecting a small perturbation yet receiving a large one). Such interaction would highlight that respective outcome’s sensitivity for prediction errors.

### Subjective and objective balance response

#### Subjective balance experience

We first assessed how expectations and perturbation intensity influenced how unstable and anxious participants felt during perturbations. We used separate generalized estimating equations (GEE) analyses to explore the effects of Expectation and Perturbation Size on both self-reported outcomes (perceived instability and anxiety). There was a significant main effect of Expectation for both perceived instability (χ^2^ = 6.05, *p* = 0.014, *d* = 0.80) and anxiety (χ^2^ = 10.38, *p* < 0.001, *d* = 1.15). There was no effect of Perturbation Size (*p*s > 0.260, *d*s <0.20), nor a significant Expectation×Perturbation Size interaction (*p*s > 0.550, *d*s <0.22). This indicates that both perceived instability and self-reported anxiety were shaped by expectation alone, with significantly greater perceived instability and anxiety levels experienced during Big-Expectation trials, irrespective of the size of the perturbation received (Figure 1B; see Supplementary Figure S1.1 for within-subject changes (connected individual datapoints) as a function of expectation and perturbation size).

#### Objective balance response (peak backwards COP velocity)

Next, we used a GEE to assess *objective* balance responses to the perturbation. In contrast to the self-reported outcomes, there was a main effect of Perturbation Size only with respect to peak backwards COP velocity (χ^2^ = 164.02, *p* < 0.001, *d* = 0.96), but no effect of either Expectation (χ^2^ = 2.35, *p* = 0.126, *d* = 0.47) nor an interaction effect (χ^2^ = 0.04, *p* = 0.835, *d* = 0.02). Consequently, the behavioural response to the perturbation was shaped by perturbation size alone, with significantly greater peak backwards COP velocity when receiving Big versus Small perturbations, irrespective of expectation (Figure 1C and 1D). This interpretation is further supported by within-subject change figures (see Supplementary Figure S1.2) which reveal an absence of consistent influence of expectation on this outcome.

### Pre-perturbation anticipatory beta activity

With respect to the EEG analysis, we first explored how expectations shaped anticipatory (pre-perturbation) EEG activity in the 1s window prior to perturbation onset. Due to the transient (or, “burst-like” (32)) nature of beta activity, we analysed the characteristics of individual beta burst-like events (15-29 Hz), rather than averaged pre-stimulus activity. Specifically, we analysed beta event rate (per/s), normalised power of beta events (normalised in factors of median (FOM)), and timing of the most recent beta event (with respect to perturbation onset; with more negative values indicating that the last beta event occurred further from perturbation onset) (25,26).

Channel-wise comparisons revealed statistically fewer pre-perturbation beta events during Big-Expectation trials across the frontal cortex – with this difference reaching statistical significance for electrode FC5 (*t* = 3.42, *p* = 0.0027, *d* = 0.75; Figure 2, top row). The mean power of pre-perturbation beta events was also significantly reduced during Big-Expectation trials across electrode Pz (*t* = 3.63, *p* = 0.0017, *d* = 0.79; Figure 2, bottom row). Both these differences remained statistically significant when corrected for multiple comparisons (*p* < 0.05). As illustrated in Supplementary Figure S2.1, the timing of the final beta event (prior to perturbation onset) was not significantly different between Small-Expectation and Big-Expectation trials across any electrodes when correcting for multiple comparisons.

**Figure 2.**
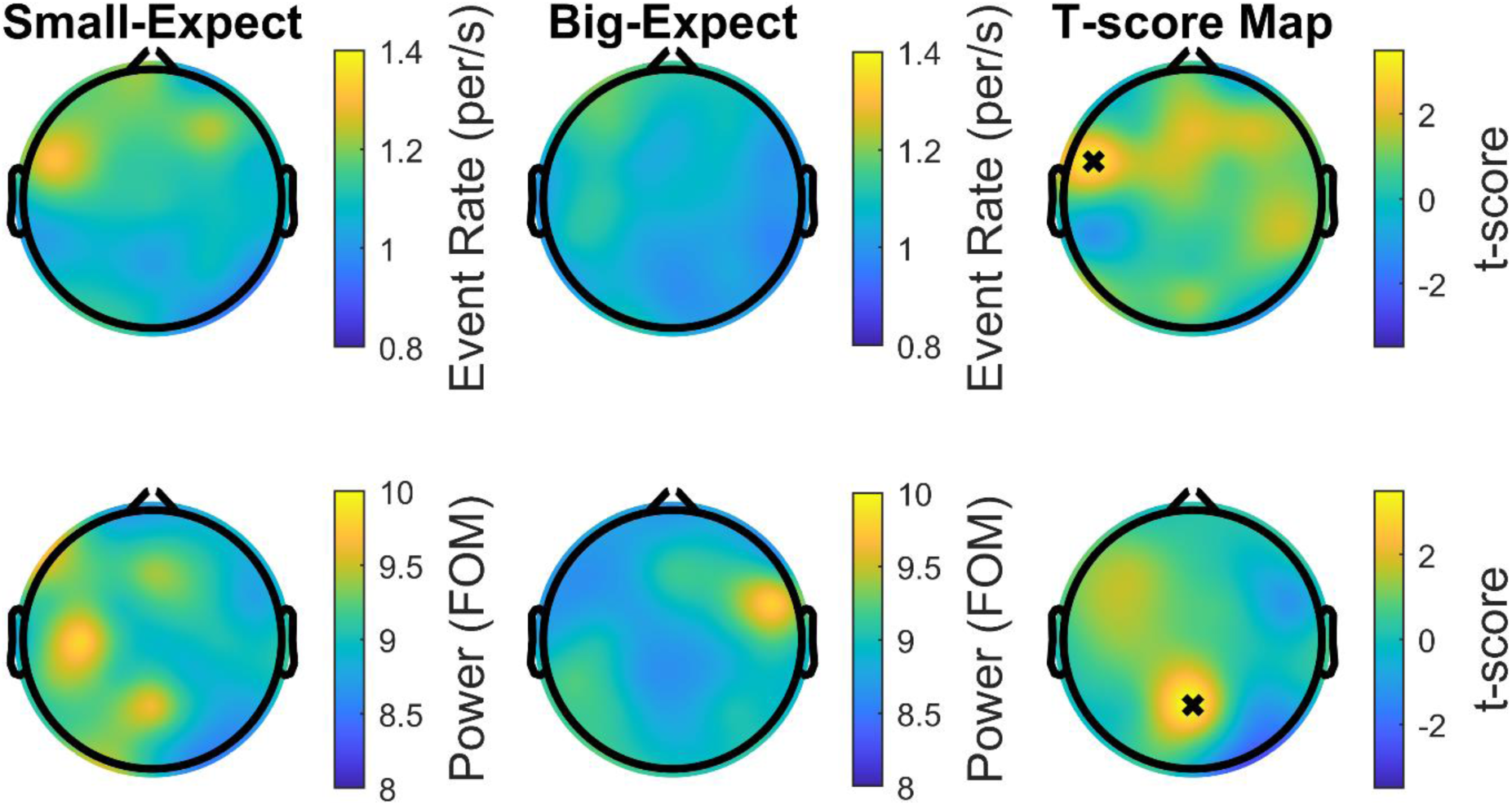
Topographical representation of beta event rate (per/s; top row) and beta mean event power (bottom row) during the one-second prior to perturbation onset. Beta mean event power is presented normalised in factors of median (FOM). X-markers indicate statistically significant differences (p<0.05, corrected for multiple comparisons).

To explore how these changes in anticipatory beta may contribute to expectation-induced perceptual biases, we examined associations between these outcomes and self-reported perceived instability. Neither frontal beta event rate (electrode FC5) nor centro-parietal beta event amplitude (electrode Pz) significantly correlated with perceived instability during either Small-Expectation (event rate, *r* = -0.39, *p_BON_* = 0.162; event power, *r* = -0.09, *p_BON_* = 1.00) or Big-Expectation trials (event rate, *r* = -0.41, *p_BON_* = 0.136; event power, *r* = 0.24, *p_BON_* = 0.600).

In summary, expectation led to clear modulations in anticipatory EEG activity, with reduced beta activity (both event rate and power of events) observed in trials when expecting bigger perturbations. However, these changes did not appear to directly relate to expectation-induced perceptual biases.

### Post-perturbation cortical outcomes

We next analysed post-perturbation cortical activity. Our analyses revealed that the balance N1 scaled to Perturbation Size (χ^2^ = 44.39, *p* < 0.001, *d* = 0.63), but not Expectation (main effect: χ^2^ = 0.35, *p* = 0.553, *d* = 0.18) with no significant interaction (χ^2^ = 0.857, *p* = 0.354, *d* = 0.24). As illustrated in Figure 3C, the balance N1 response was significantly larger (i.e. greater negativity) when receiving Big versus Small perturbations, irrespective of expectation. In contrast, the within-subject change figures (Supplementary Figure S1.3) reveal an absence of consistent or systematic influence of expectation on this outcome. This is therefore in line with the objective balance outcomes, suggesting that the balance N1 is shaped by the received, rather than expected, sensory signals. A detailed visualisation of individual cortical N1 waveforms and topographical morphologies are presented in Supplementary Material 3.

**Figure 3.**
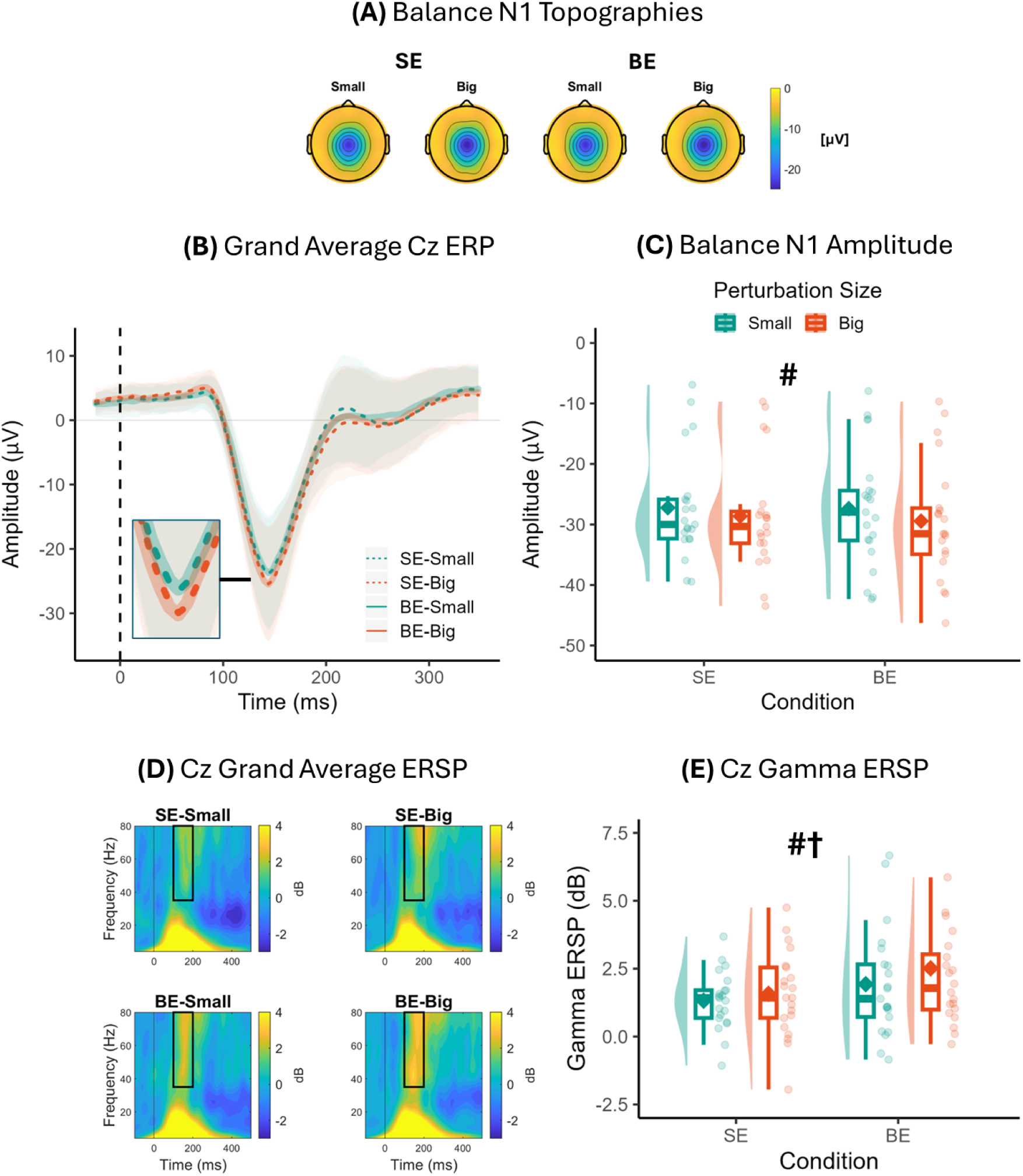
**A)** Post-perturbation cortical responses during the four trial types, including grand average balance N1 scalp topographies. **B)** Grand average (±SD) ERP of channel Cz. **C)** Balance N1 amplitudes. **D)** Grand average ERSP plots of channel Cz across each trial type. **E)** Mean Cz gamma (35-80 Hz) ERSP. Note, SE reflects Small-Expectation trials whilst BE reflects Big-Expectation trials (e.g., ‘SE-Small’ indicates expecting a small perturbation and receiving this). Timepoint zero on 3B and 3D indicates perturbation onset. Condition means for boxplots on 3C and 3E are denoted with ◊. Significant main effects of Expectation (†) and Perturbation Size (#) are denoted on the Figures where relevant. Please see the Supplementary Material 1 for boxplots showing within-subject changes from Figure 3C (Supplementary Figure S1.3) and 3E (Supplementary Figure S1.4) as a function of expectation and perturbation size.

We next analysed evoked sensorimotor (channel Cz) gamma activity, focusing on a time-window 100-200ms after perturbation onset (solid black box, Figure 3D). In contrast to the balance N1 which scaled to perturbation size only, post-perturbation gamma activity independently scaled to both Expectation (χ^2^ = 5.28, *p* = 0.022, *d* = 0.49) and Perturbation Size (χ^2^ = 6.77, *p* = 0.009, *d* = 0.27). The interaction effect was not significant (χ^2^ = 1.41, *p* = 0.235, *d* = 0.21). As illustrated in Figure 3E, post-perturbation gamma activity was both higher during Big-Expectation compared to Small-Expectation trials (irrespective of the size of the subequent perturbation), and when receiving Big versus Small perturbations (independent of expectation; see Supplementary Figure S1.4 demonstrating consistent within-subject changes as a function of both expectation and perturbation size). Interestingly, we observed a strong, significant correlation between post-perturbation gamma and self-reported instability during Big-Expectation trials (*r* = 0.69, *p_BON_* = 0.001; see Supplementary Figure S2.2). However, the correlation between these two variables during Small-Expectation trials did not reach statistical significance following Bonferroni correction (*r* = 0.49, *p_BON_* = 0.051).

In summary, our results revealed that post-perturbation evoked EEG activity is independently shaped by both expectation and sensory signals. Whilst the balance N1 scaled to the size of the perturbation alone (with no influence of expectation), post-perturbation gamma activity was influenced independently by both expectation and perturbation size. Interestingly, post-perturbation gamma was also strongly associated with self-reported perceived instability during Big-Expectation trials. Contrary to our predictions, there was no evidence of either the balance N1 or post-perturbation gamma being shaped by prediction errors, as no significant interaction effects were observed.

## Discussion

Our findings provide direct evidence that top-down expectations can bias how sensory signals are processed at both the perceptual and neural level during reactive balance control. Participants reported feeling more unstable when expecting greater postural instability—even when sensory input (and actual behavioural responses) remained constant. These perceptual changes were accompanied by distinct EEG signatures: expecting greater imbalance reduced anticipatory beta-band activity and increased post-perturbation gamma responses (with the latter being strongly associated with expectation-induced perceptual biases). In contrast, the balance N1 evoked response, as well as behavioural responses themselves, were influenced by perturbation magnitude only. Notably, none of the a priori EEG measures showed evidence of strong sensitivity to prediction error, suggesting that mismatches between expected and actual perturbations did not appear to drive the dominant short-latency cortical responses observed. Instead, it appears that two dissociable processes may exist during reactive balance control: one that is sensitive to top-down expectations and another that is driven by sensory input, with limited evidence for sensitivity to mismatch or prediction error within the early temporal window examined here. We propose that such dual system allows for perceptual stability whilst ensuring that behaviour can be flexibly tuned to rapidly changing environmental contexts.

### Top-Down Modulation of Beta and Gamma Activity

Our results highlight two distinct, expectation-sensitive EEG markers: anticipatory beta and post-perturbation gamma. Reduced beta power in sensory regions prior to perturbation onset was observed when participants expected greater instability. This mirrors findings from pain research, where high-expectancy of painful stimuli leads to reduced pre-stimulus beta and increased sensory sensitivity (8). Beta oscillations are commonly interpreted as reflecting cortical inhibition (25,26,33); thus, reductions may reflect a hypervigilant state that primes the system for heightened responsiveness to expected input (34). In the context of balance, this may result in amplified neural processing of (and exaggerated behavioural responses to) even mild postural disturbances, as commonly seen in various clinical balance disorders (19,35).

The inhibitory role of cortical beta is not restricted to sensory processing, as it also plays a key role in *motor* inhibition, particularly in frontal regions. For example, successful inhibition of an upper limb action has been linked to increased frontal beta events (35,36). The demands of the present study required participants to rapidly initiate rather than *inhibit* a motor action to contend with the postural perturbation (i.e. a compensatory postural reaction). As we observed significant reductions in frontal beta events when participants expected the upcoming perturbation to be more destabilising, we propose that this may reflect a top-down strategy to facilitate rapid motor initiation. Future work could directly test this hypothesis by combining EEG with measures of neuromuscular responses under varying expectation conditions. Whilst this interpretation should be considered in light of the relatively modest reductions in beta-event rate that we observed (mean decrease from 1.31 (*SD* = 0.38) to 1.09 (*SD* = 0.28) events per/s; *d* = 0.75), prior work has reported that even relatively small changes in beta-event rate (36) and/or power (37,38) can meaningfully modulate subsequent perception and behaviour. Together, reductions in sensory beta power and frontal beta events suggest complimentary mechanisms by which expectations prime the sensorimotor system for anticipated imbalance.

Post-perturbation sensorimotor gamma activity (100–200ms) increased both when participants expected greater instability, and when they experienced it. Although we initially hypothesised that gamma would track prediction error (see (30)), our findings suggest that post-stimulus gamma responses may instead reflect perceived sensory salience more broadly: stimuli that demand greater processing (or which are expected to) elicited stronger gamma activity. This aligns with evidence that gamma oscillations support sensory gain and perceptual amplification (39,40). In our study, gamma responses were independently shaped by expectation, and were strongly associated with subjective reports of perceived instability during Big-Expect trials (*r* = 0.69, *p_BON_* = 0.001). These findings provide novel evidence that instability-based gamma may contribute to the neural processes underpinning balance perception.

### Prediction errors: Hard to predict?

Contrary to our hypotheses, no a priori EEG markers, including the balance N1, tracked the mismatch between expected and received perturbations (i.e., prediction error). Rather, the N1 scaled to the size of the perturbation received, supporting previous work describing larger balance N1 responses during more destabilising events (27,41). Although some have proposed that the balance N1 reflects an ‘error detection’ mechanism (42), our findings support the alternative view that the N1 reflects a cortical (but subconscious) mechanism employed to detect and behaviourally respond to instability, rather than signalling error per se. Supporting this interpretation are findings that larger N1 responses are recorded during trials in which participants must take a compensatory step to regain balance following a perturbation (41,43). While prediction errors (likely originating from subcortical areas such as the cerebellum (44)) are known to influence longer-term balance behaviour (e.g. locomotor adaption (21,22)), our findings suggest these exert little influence over either conscious balance perception or early cortical responses likely to shape immediate behavioural responses. However, by unpredictably interleaving expectation-violating trials with expected trials (as in the present work and previously (7,8)), the design prioritises immediate perceptual and neural biases, precluding assessment of longer-term adaptation processes. Another important point relates to the reactive (rather than volitional) nature of the present motor task. Thus, although our findings suggest dissociable systems governing short-latency reactive balance responses, they may not generalise to volitional balance behaviours such as walking, where internal forward models are rapidly updated in response to prediction errors (21,22).

Given the typical latencies of the cortical pathways associated with reactive balance responses to comparatively small perturbations such as those used here (45), we focused our analyses on early sensorimotor cortical responses (i.e., ≤200ms). However, it is possible that other neural markers outside this temporal window might instead reflect error processing during balance. For instance, Jalilpour and Muller-Putz (46) found that balance perturbations characterised by a prediction error (participants tilted in opposite direction as expected) were only distinguished from trials that matched expectations by cortical rhythms occurring following the N1. Specifically, they identified post-N1 alpha suppression to be a key marker of error processing (a response often termed ‘error-related alpha suppression’ (46)). In-line with this, our exploratory whole-scalp analyses (see Supplementary Figure 2.2) revealed a similar suppression of alpha activity (200–400ms) over parietal-occipital electrodes following unexpectedly *large* perturbations (i.e., big perturbations during the Small-Expect block), but not following unexpectedly *small* ones. These findings suggest that longer-latency post-N1 alpha suppression may serve as a neural marker of balance error processing (particularly when these errors could have negative real-world consequences, e.g., instability was larger than predicted), potentially influencing later behavioural responses beyond the initial backward instability examined here. Because our primary analysis focused on early neural and behavioural outcomes that are ecologically relevant for preventing a fall, our conclusions are accordingly restricted. Future work should therefore systematically investigate how expectation and prediction errors affect longer-latency responses to instability.

### Implications for Balance Disorders and Predictive Coding

That expectations alone can both distort balance perception and modulate cortical activity has important implications for clinical conditions marked by subjective instability, including functional dizziness (47) and age-related balance disorders (19,48)). Identifying EEG markers of expectation-driven perceptual distortion, particularly anticipatory beta and post-perturbation gamma may offer new diagnostic or monitoring tools for these conditions. Currently, there are none.

More broadly, our findings extend perceptual predictive coding frameworks into the motor domain of balance regulation. Unlike more traditional sensory systems, balance involves complex multisensory integration and rapid subcortical feedback mechanisms (10,11). Yet even here, top-down expectations shape perception and modulate cortical responses. Importantly, unlike pure uni-sensory processes (4,49,50), these effects do not appear to rely on prediction errors, but rather on direct modulation of neural activity linked to motor preparation and sensory salience.

### Conclusions

In sum, our results demonstrate that early cortical processing of imbalance integrates both sensory input and top-down expectations, but not their mismatch. Notably, subjective balance perception was shaped by expectations alone. These findings identify anticipatory beta and post-perturbation gamma as specific EEG markers that may mediate perceptual distortions in balance and motor control more broadly. This helps refine predictive coding frameworks beyond purely sensory domains but also points to potential neural targets for diagnosing or treating balance disorders rooted in expectation-based distortions (51).

## Methods

### Participants

Twenty-one neurotypical adults participated in the experiment (11 females, 10 males; M ± SD age = 25.0 ± 4.9 years; height = 173.0 ± 11.0 cm; weight = 73.7 ± 10.9 kg). Participants were free from neurological disease and had no prior experience of dizziness or balance problems. The experiment was approved by the Manchester Metropolitan University ethics committee. Sample size estimates were based on the self-report measure of perceived instability, our primary indicator of expectation manipulation. Prior studies have reported large to very large effects for similar manipulations on perceptual outcomes during balance (52,53). Power analysis for a 2×2 repeated-measures, within-subjects design (f = 0.40, α = 0.05, power = 0.80, r = 0.70) indicated that 14 participants were sufficient to detect similarly large effects.

### Protocol

Perturbations were delivered by a moveable platform (80×60 cm with an embedded force plate recording at 1,000 Hz; Type 9281B, Kistler Instrument). The platform was driven by an electromagnetic actuator and controlled through custom written software (LabVIEW v19 SP1, National Instruments) via DAQ card (USB-6210, National Instruments). Participants stood on the force plate, with feet shoulder width apart and hands on hips. Foot positioning was marked to ensure consistency. During perturbations, participants were instructed to fixate on a cross marked on the wall at eye level, 4 m away. Participants wore noise-cancelling headphones to minimise any anticipatory audio cues.

### Expectation manipulation

Participants experienced two blocks of 60 discrete sine wave perturbations (7-15s random delay between each perturbation), each consisting of an initial forward translation of the support surface before reversing in direction and completing the sine wave to return to the original position. Both blocks contained a combination of “Small” (maximum forward displacement = 57.0mm; peak acceleration = 1,472mm/s²; peak acceleration latency = 70ms; see Figure 1) and “Big” perturbations (maximum forward displacement = 73.3mm, peak acceleration = 1,965mm/s²; peak acceleration latency = 60ms; see Figure 1A). Prior to completing the two experimental blocks of trials (described below), participants received 5 small and 5 big perturbations (order randomised across participants) to enhance the formation of their expectations.

In the Small-Expectation (SE) block of trials, participants were informed they would encounter small perturbations throughout the block of 60 trials. This block commenced with ten consecutive small perturbations, reinforcing their expectation. The remaining 50 perturbations comprised a predetermined sequence of 35 small and 15 big (expectation-violating) perturbations. These expectancy-violating perturbations were introduced on average every third trial and were never presented in succession. Such interleaved approach (in line with prior work (7,8)) allowed us to probe how expectations shape perception and neural processing under dynamic and uncertain conditions. The Big-Expectation (BE) block of trials followed the same structure but with the perturbation sizes reversed. The sequence of perturbations was fixed and identical for all participants, ensuring consistency across the cohort, and the presentation of the two blocks of trials was counterbalanced across participants.

To ensure balanced counts across trial-types, we compared 15 Small perturbations from the Small-Expectation block to 15 Small perturbations from the Big-Expectation block, and 15 Big perturbations from the Big-Expectation block to 15 Big perturbations from the Small-Expectation block. To ensure a consistent and unbiased selection, the 15 trials (out of the available 45) selected from the expectancy-aligned condition (i.e., small perturbations during the Small-Expectation condition) were defined as those immediately preceding a trial that violated expectancy (see Supplementary Figure 4 for a schematic example of how these different trial types were selected). This selection criterion was applied for two key reasons: (a) the first 10 trials of each block were excluded to avoid confounding effects of early adaptation, and (b) the selected trials were always maximally distant from a preceding expectancy-violating perturbation, reducing potential carryover effects.

Both perturbation sizes (small and big) were designed to challenge postural stability without necessitating a corrective stepping response. To prevent fatigue, participants received a 5–10 minute break between each block of 60 perturbations.

### Self-reported outcomes

After certain trials, participants completed self-reported questionnaires assessing their perceived instability and anxiety during the preceding trial (17). For perceived instability, participants were asked “how unstable did you feel during the trial?” and responded via a visual analogue scale (0 = “completely steady”, 10 = “so unsteady that I would fall”). Participants were then asked to rate the level of anxiety they experienced in the preceding trial via a visual analogue scale that ranged from 0 “not at all anxious” to 10 “extremely anxious” (17,27). These questionnaires were completed after two random big and small perturbations, in each expectation block.

### EEG recording and analyses

EEG signals were recorded at 1000 Hz from 29 active electrodes (eego sports, ANT Neuro, Hengelo, Netherlands) positioned according to the extended 10–20 international system. Signals were band-pass filtered (1–80 Hz), epoched (−2 to +2s relative to perturbation onset) and averaged referenced. Independent component analysis (ICA; RunICA informax) were then conduced to manually reject non-neural activity (average of 23.5 ± 1.9 components retained per participant).

Pre-perturbation cortical beta (15 – 29 Hz) activity was analysed across the 1s pre-perturbation window, given the clear associations between this outcome and both subsequent stimulus perception (25,26,32) and behavioural response (36,37). Following the methods of Shin et al. (26), we measured three key outcome measures using the open-source ‘SpectralEvents Toolbox’ within MATLAB (54): beta event rate (per/s), normalised power of beta events (normalised in factors of median (FOM)), and timing of the most recent beta event (with respect to perturbation onset; ranging from -1 to 0 s). Please see Supplementary Materials 4 for further details about the analyses employed.

For post-perturbation data, we calculated the ‘balance N1’ for each trial as the largest negative peak occurring at electrode Cz 50-200ms after perturbation (following baseline correction [-1000 to -250ms]; 27,28). We also calculated Event-Related Spectral Power (ERSP) for electrode Cz, using complex Morlet wavelets. We used 77 frequencies linearly spaced between 4 and 80 Hz, with wavelets logarithmically spaced from 3 to 8 cycles. Data were baseline normalised to average activity from -500 to -200ms prior to perturbation onset. We chose to focus on gamma based on the well-documented role of cortical gamma as a robust prediction error signal across tasks and sensory modalities (30,31). We limited our analysis to 35-80Hz (to avoid also capturing any beta activity) 100-200ms post-perturbation onset. This window of interest was selected based on condition-average inspection that revealed gamma-band activity to be strongest within this boundary (55).

### Postural response to perturbations

We used custom MATLAB scripts to determine the peak backwards velocity of anterior-posterior (AP) centre of pressure (COP) data in response to the initial forward portion of the perturbation. Peak backwards COP velocity was selected as our outcome variable as it is a direction-specific response to the initial forward perturbation; greater backwards CoP velocity generally indicates greater instability and higher risk of falling (56,57). First, we selected and low-pass filtered (5Hz, 2nd order bidirectional Butterworth filter) a 3-second AP-COP trace that spanned 2000ms pre-perturbation and 1000ms post-perturbation. We then corrected this trace for offset using the estimated average AP COP displacement during the ‘baseline’ period (based on the 1100-100ms pre-perturbation window). Peak velocity of the postural response to the perturbation was then identified as the first negative peak in the derivative of the AP-COP trace in the initial forward portion of the perturbation (Figure 1). By default, the initial negative peak was selected unless a subsequent peak was of >50% greater magnitude than the earlier peak (27).

### Statistical analyses

#### Subjective and objective balance outcomes

Subjective (perceived instability and anxiety) and objective balance outcomes (AP COP velocity) were analysed using separate generalized estimating equations (GEE) analyses to explore the effects of Expectation (SE and BE) and Perturbation Size (Small and Big). We chose an exchangeable working correlation matrix to define dependency amongst measurements. Significant main and interaction effects were followed up with Bonferroni-corrected post-hoc pairwise comparisons. Threshold for significance was set at α = 0.05.

#### Pre-perturbation analyses

To assess the influence of expectation on preparatory cortical activity, trial-level beta outcomes were averaged across all trials within an expectation block. This approach enabled us to specifically examine how Expectation (i.e., Small-Expectation vs. Big-Expectation) modulated preparatory neural activity prior to perturbation onset. Given this activity occurred *before* the delivery of an expectancy-violating trial, averaging within blocks provided the most appropriate means of capturing expectation-driven cortical dynamics (and served to maximise analytical power). To explore expectation-driven topographical differences between experimental blocks in pre-perturbation beta activity, we performed channel-wise paired samples t-tests. To control for inflated Type I error due to multiple comparisons across EEG channels, we employed a non-parametric permutation-based “maximum statistic” method (55).

#### Post-perturbation outcomes

We used generalized estimating equations (GEE) analyses to explore the effects of Expectation (SE and BE) and Perturbation Size (Small and Big) on both the balance N1 and post-perturbation gamma. This approach was deemed the most appropriate for our time-frequency analyses given the difficulties associated with applying permutation testing to factorial designs as in the present study (e.g., Expectation x Perturbation Size; see (55)). As with the balance outcomes described above, we chose an exchangeable working correlation matrix. Significant main and interaction effects were followed up with Bonferroni-corrected post-hoc pairwise comparisons. Threshold for significance was set at α = 0.05.

#### Correlations

To further evaluate how EEG changes may contribute to expectation-induced perceptual biases, we conducted Pearson correlations examining associations between self-reported perceived instability and the EEG measures that were sensitive to expectations (anticipatory frontal beta event rate, centro-parietal beta event power, and post-perturbation gamma). Separate correlations were performed for Small-Expectation and Big-Expectation trials (expectation confirming and violating trials averaged together given the main effect of expectation, but lack of interaction effect). Bonferroni corrections were applied to control for multiple comparisons, and the significance threshold was set at α = 0.05.

#### Effect size estimates

We report standardised effect sizes using Cohen’s d. For GEE analyses, we operationalised these as standardised marginal effects, which were obtained by dividing GEE estimated marginal contrasts by the between-participant standard deviation for the respective outcome measure.

## Supplementary material

### 1. Extended data visualisation

The following figures are intended to illustrate the within-participant change scores that underpin the generalised estimating equation (GEE) analyses reported in the main manuscript. Each figure contains four panels showing the effect of Perturbation Size for the Small-Expectation (SE [Panels A]) and Big-Expectation (BE [Panels B]) conditions, and the effect of Expectation for both the Small (Panels C) and Big (Panels D) perturbations. Each panel contains raincloud-style plots that display individual data points connected across repeated measures with lines. Half-violin density plots and boxplots summarise data distributions, whilst diamond symbols represent the mean. The right-hand section of each panel displays participant-level difference scores (i.e., Δ = Big – Small, or Δ = BE - SE) for visualisation of within-participant effects.

**Figure S1.1.**
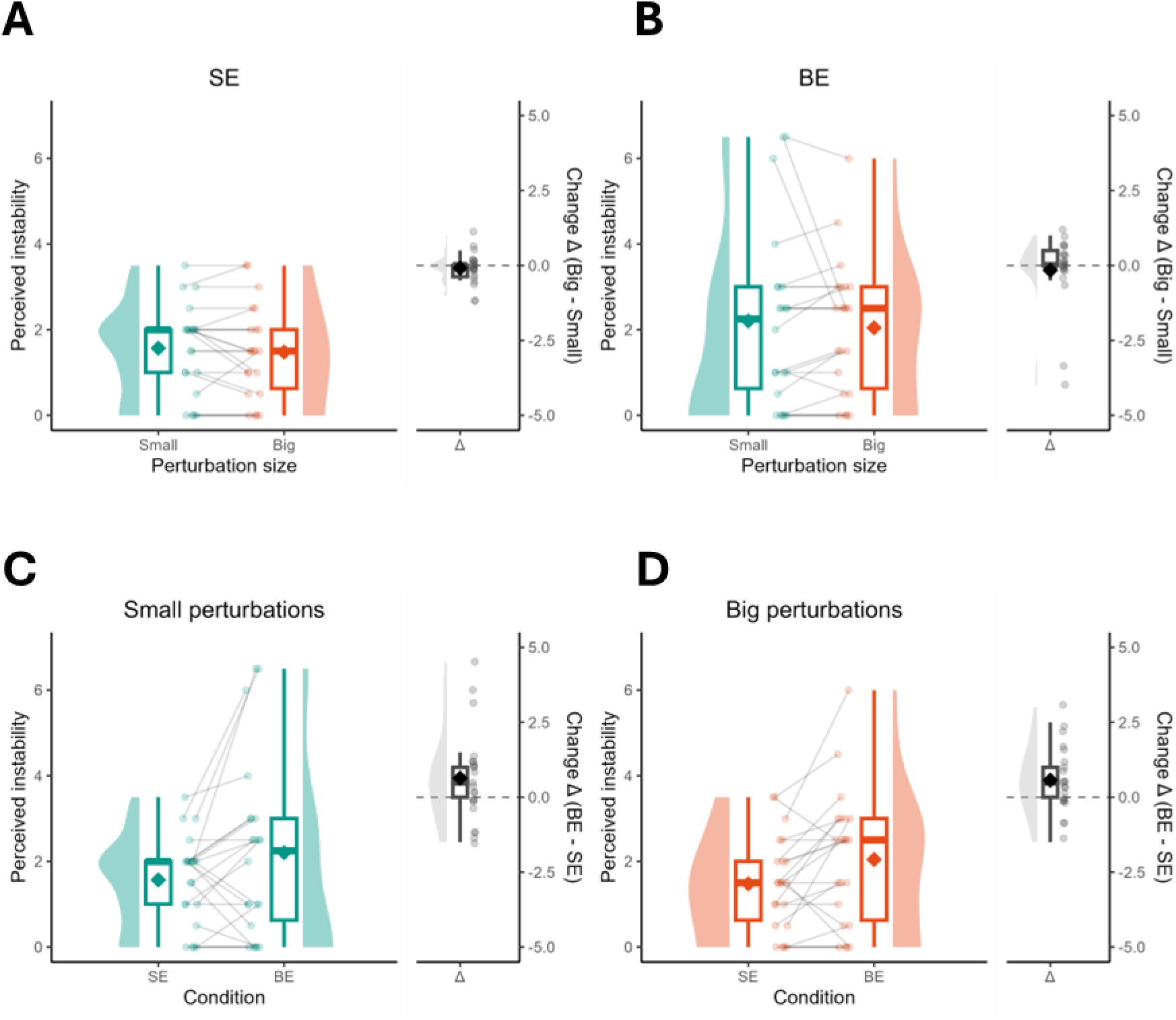
Data visualisation of perceived instability to highlight the effect of Perturbation Size (panel A [during SE trials] and panel B [during BE trials]) and Expectation (panel C [during Small perturbations] and panel D [during Big perturbations]).

**Figure S1.2.**
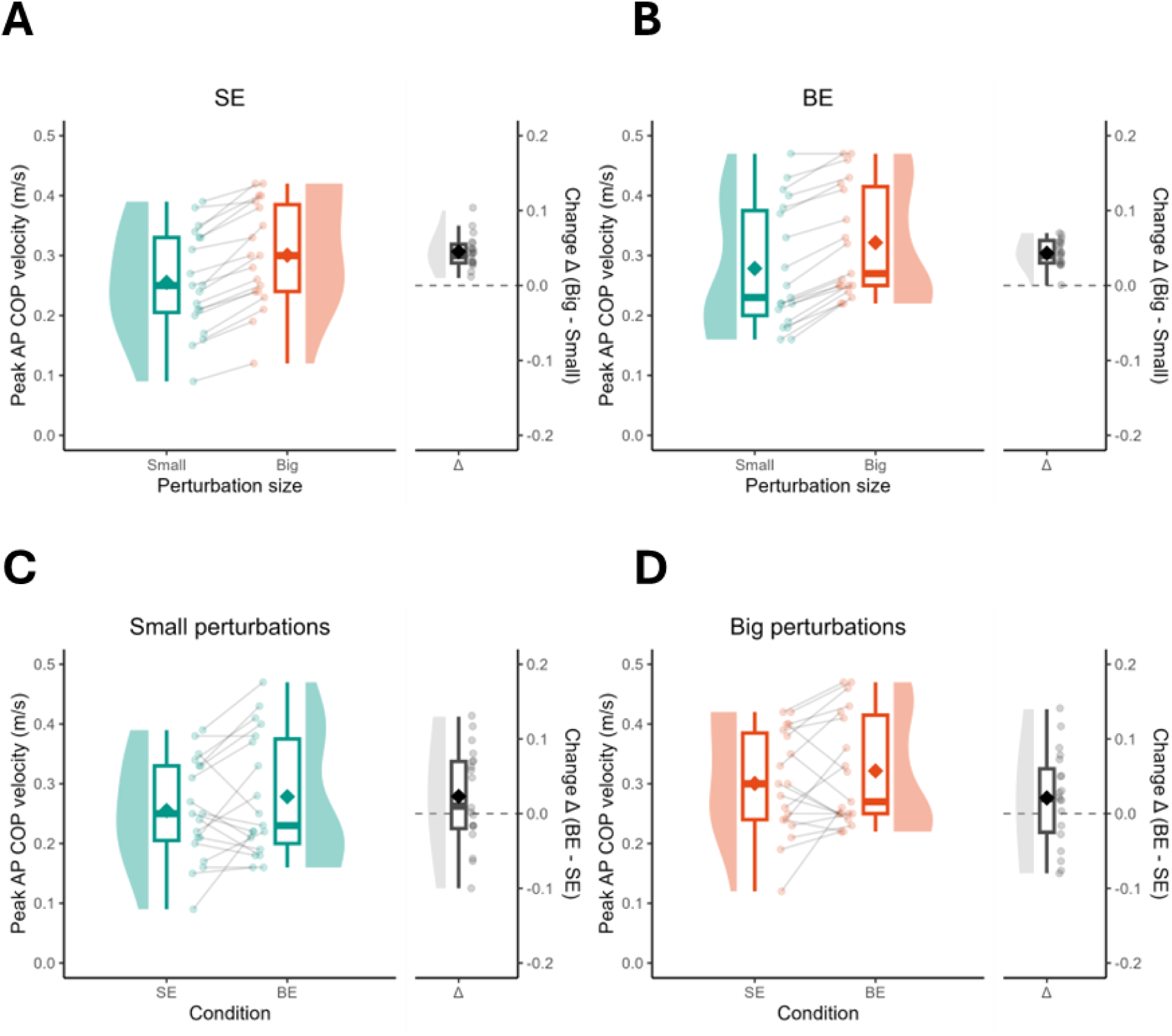
Data visualisation of peak backwards anteroposterior (AP) centre of pressure (CoP) velocity (m/s) to highlight the effect of Perturbation Size (panel A [during SE trials] and panel B [during BE trials]) and Expectation (panel C [during Small perturbations] and panel D [during Big perturbations]).

**Figure S1.3.**
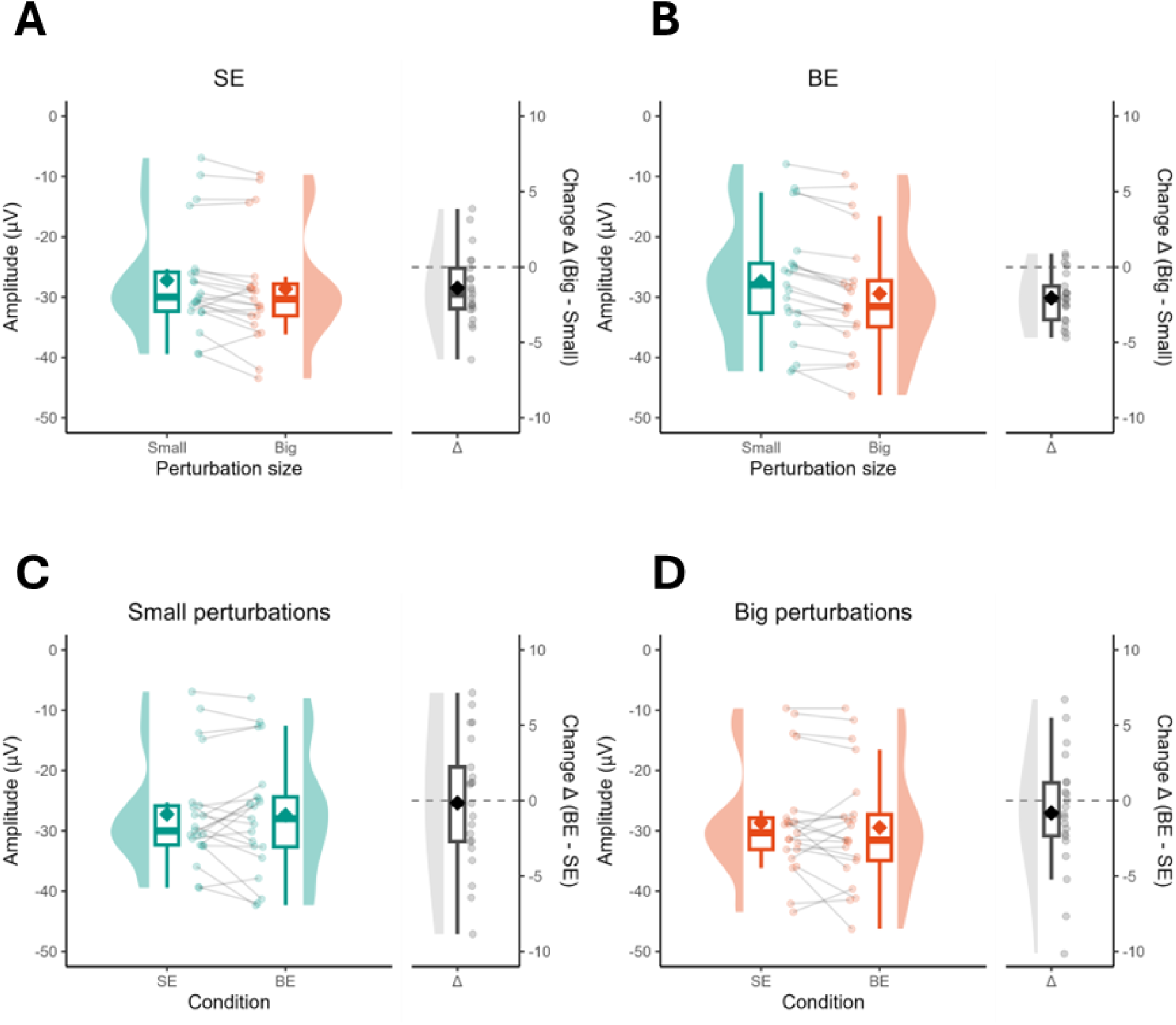
Data visualisation of cortical N1 amplitudes (µV) to highlight the effect of Perturbation Size (panel A [during SE trials] and panel B [during BE trials]) and Expectation (panel C [during Small perturbations] and panel D [during Big perturbations]).

**Figure S1.4.**
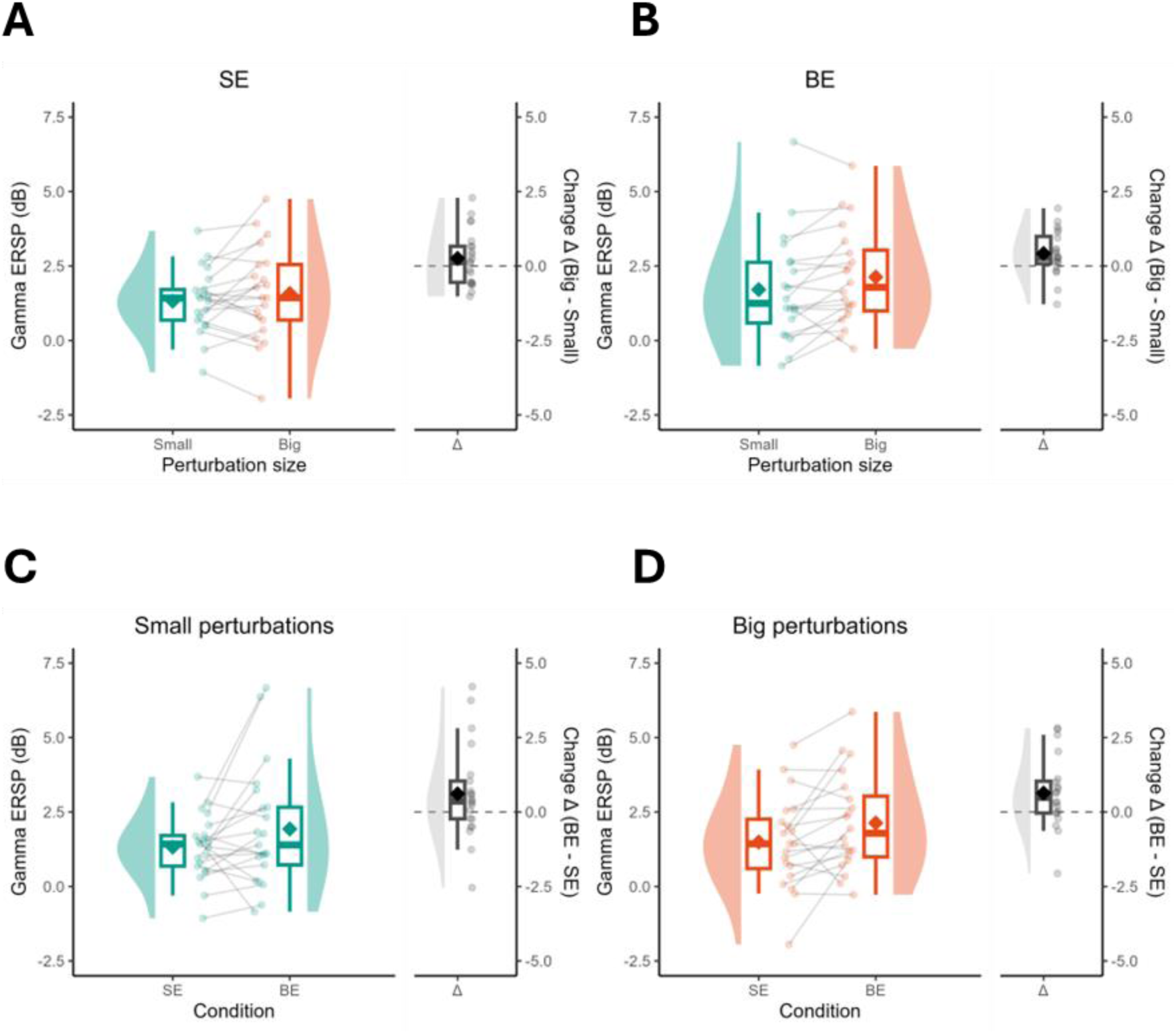
Data visualisation of post-perturbation (100-200 ms) beta event-related spectral power (ERSP; dB) to highlight the effect of Perturbation Size (panel A [during SE trials] and panel B [during BE trials]) and Expectation (panel C [during Small perturbations] and panel D [during Big perturbations]).

### 2. Additional analyses

As illustrated in Supplementary Figure S2.1, the timing of the final beta event (prior to perturbation onset) tended to also be earlier (i.e., further from perturbation onset) during Big-Expectation trials—specifically for frontal (F4 (*t* = 2.19, *p* = 0.040, *d* = 0.48) and FC5 (*t* = 2.79, *p* = 0.011, *d* = 0.61)) and central electrodes (C4; *t* = 2.20, *p* = 0.040, *d* = 0.48). However, these differences did not survive statistical correction for multiple comparisons.

**Figure S2.1.**
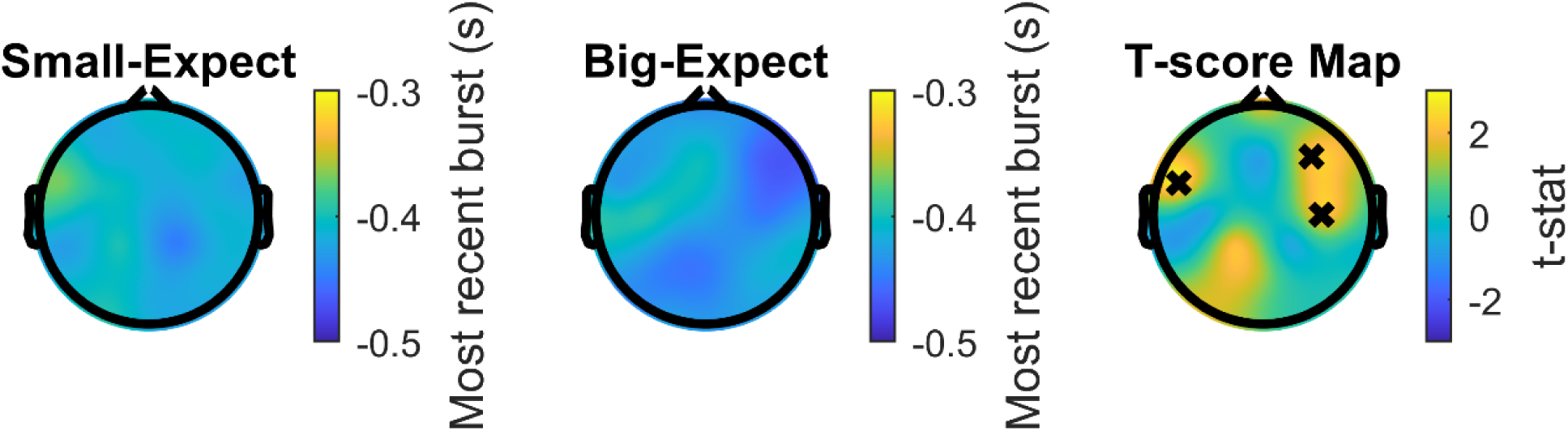
Topographical representation of timing of the final beta event, prior to perturbation onset. Note, time zero indicates the time at which the perturbation occurred (i.e., an event occurring at -0.3s thus occurs 300ms prior to perturbation onset). X-markers on the t-score map indicate statistically significant differences between conditions; however, these differences did not survive statistical correction for multiple comparisons.

**Figure S2.2.**
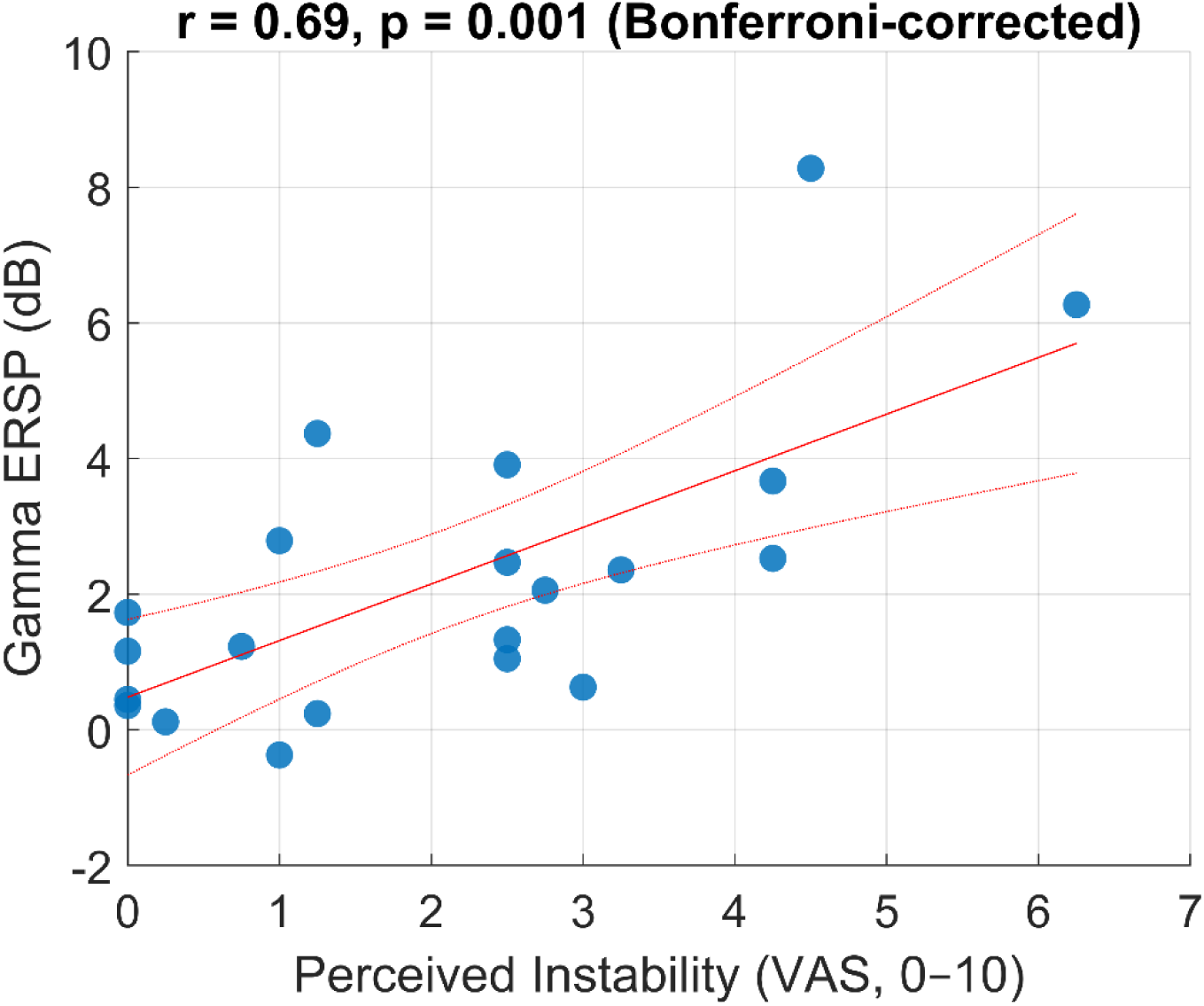
Scatter plot displaying the positive linear correlation between perceived instability and post-perturbation gamma ERSP (dB) during Big-Expectation (BE) trials.

**Figure S2.3.**
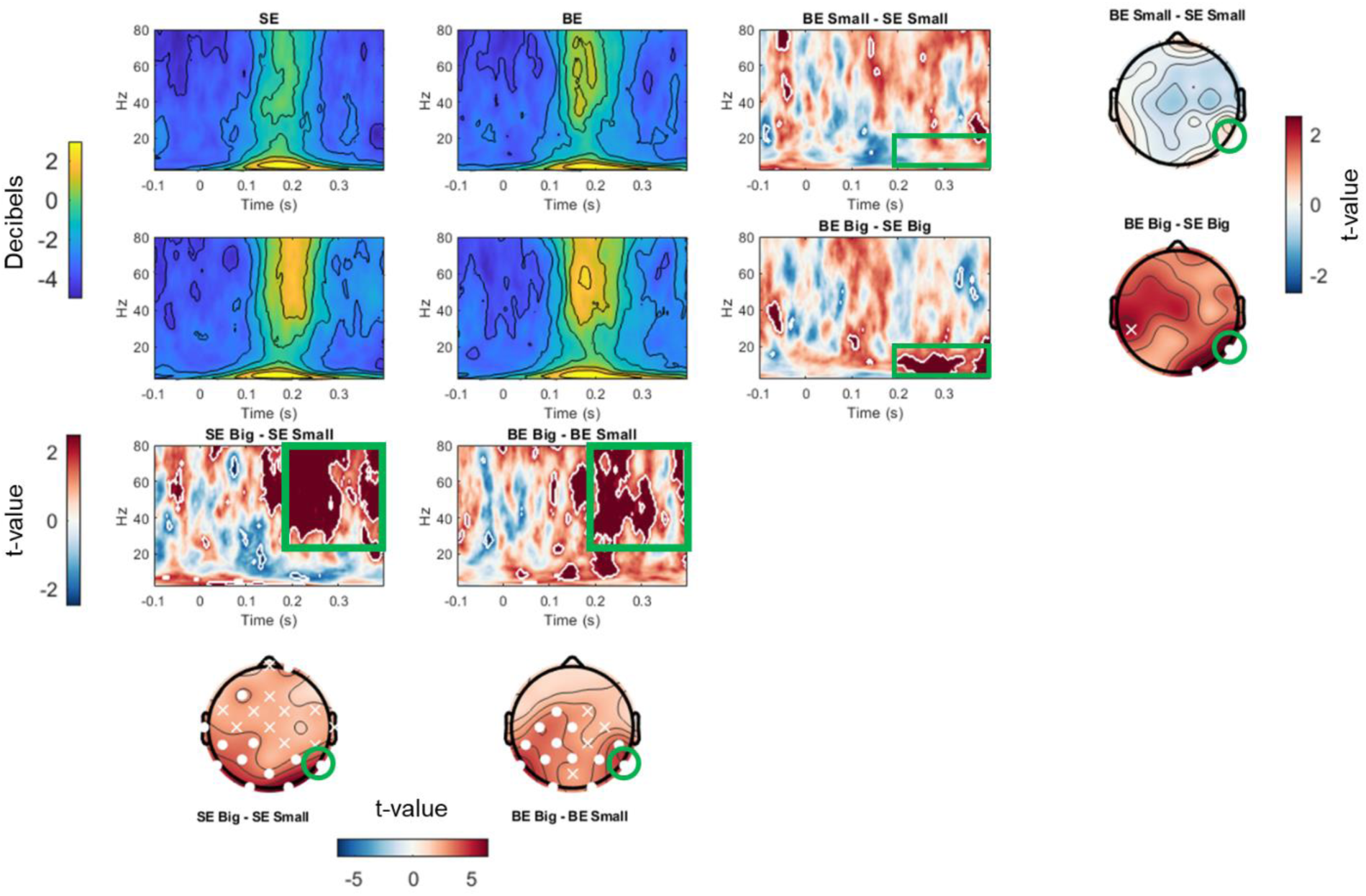
Event-related spectral power and statistical comparisons following perturbations (exploratory analysis). This figure shows time–frequency representations of grand average event-related spectral power (ERSP) at electrode P6 across all perturbation conditions. The first two columns depict ERSPs (in decibels) for each perturbation size (rows) and expectation condition (columns): Big perturbations are shown in the middle row, Small perturbations in the top row. Within each row, data are grouped by Expectation condition: Big-Expecting (BE; left column) and Small-Expecting (SE; middle column). Warmer colours (i.e., yellow) indicate increased spectral power relative to baseline. The third column displays pixel-wise t-value maps comparing ERSPs across Expectation conditions. For Small perturbations (top), ERSPs during the SE block are compared to those during the BE block. For Big perturbations (middle), SE-Big is compared to BE-Big. Red areas reflect greater power during BE blocks, while blue reflects greater power during SE blocks. White outlines indicate statistically significant clusters. Notably, a marked reduction in alpha activity (8–12 Hz) between 200–400 ms was observed when perturbations were larger than expected (i.e., SE-Big vs BE-Big). The adjacent topoplot shows the spatial distribution of this alpha difference, with significant effects (white circles) observed over parieto-occipital electrodes (P6 and O2), and sub-threshold effects marked with an “×” (e.g., CP5). The bottom row compares ERSPs between Big and Small perturbations within each Expectation condition (left = SE, right = BE). These contrasts revealed increased beta–gamma (25–80 Hz) activity from ∼150–400 ms following larger perturbations. Corresponding topographical plots show this activity was broadly distributed across posterior electrodes.

### 3. Visualisation of individual balance N1 morphology

The following supplementary figures provide complementary visualisations of the balance N1 response across Expectation and Perturbation Size. These figures are intended to illustrate individual-level variability in N1 waveform morphology, latency, and scalp topography, and to aid interpretation of the group-level analyses reported in the main manuscript. To prioritise visual clarity of within-participant patterns, several figures use participant-specific normalisation, temporal alignment, or scaling; consequently, these visualisations are descriptive and are not intended for quantitative comparison of absolute amplitudes between participants.

**Figure S3.1.**
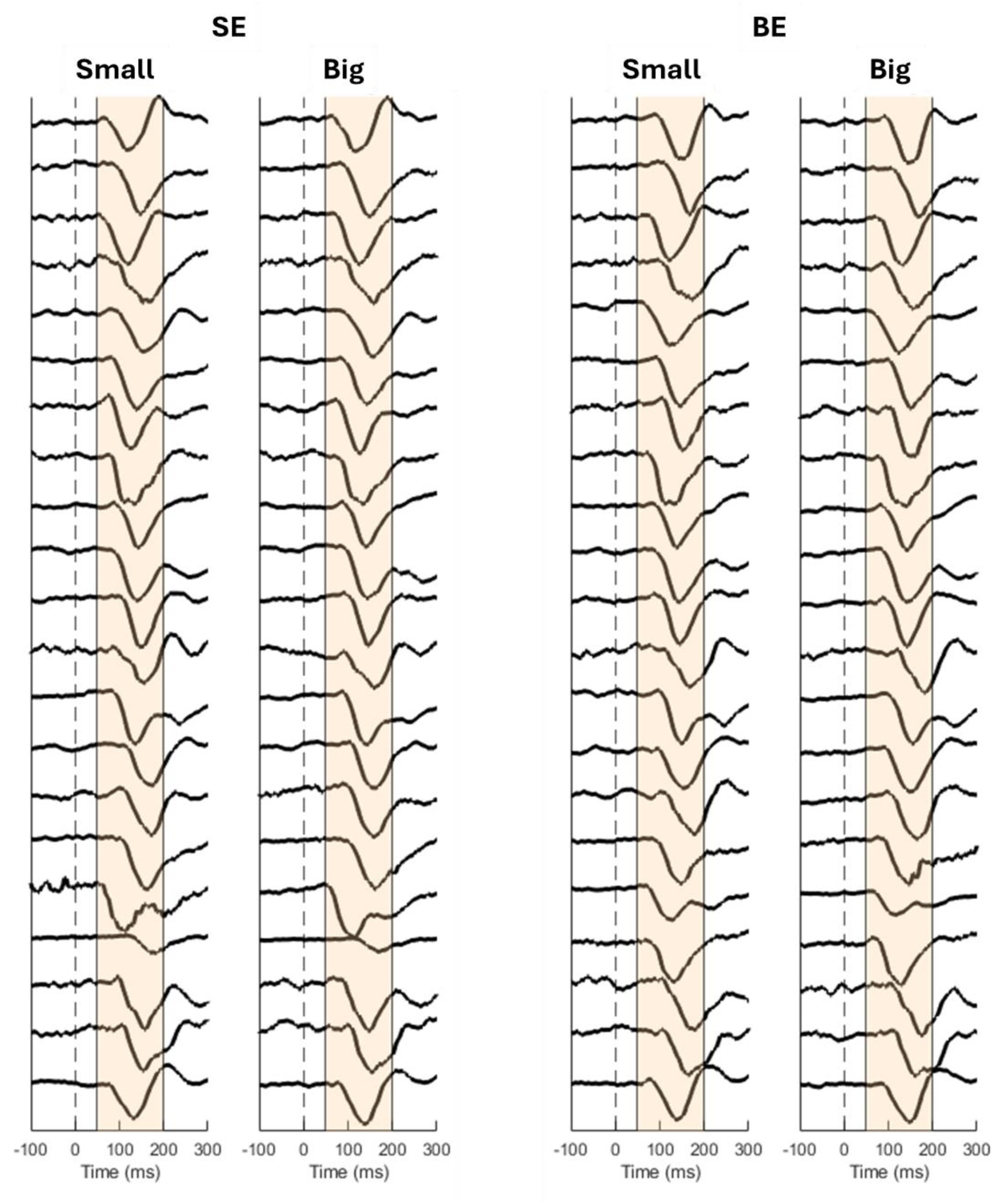
Visualisation of each participant’s grand-average balance N1 waveform at Cz for every Condition × Perturbation Size combination. The shaded yellow region marks the 50–200 ms window used to quantify peak N1 amplitude, and the vertical dashed line indicates perturbation onset (time 0). To maximise visibility of individual N1 morphology, each participant’s waveform was normalised within-participant; therefore, amplitudes are not comparable between participants.

**Figure S3.2.**
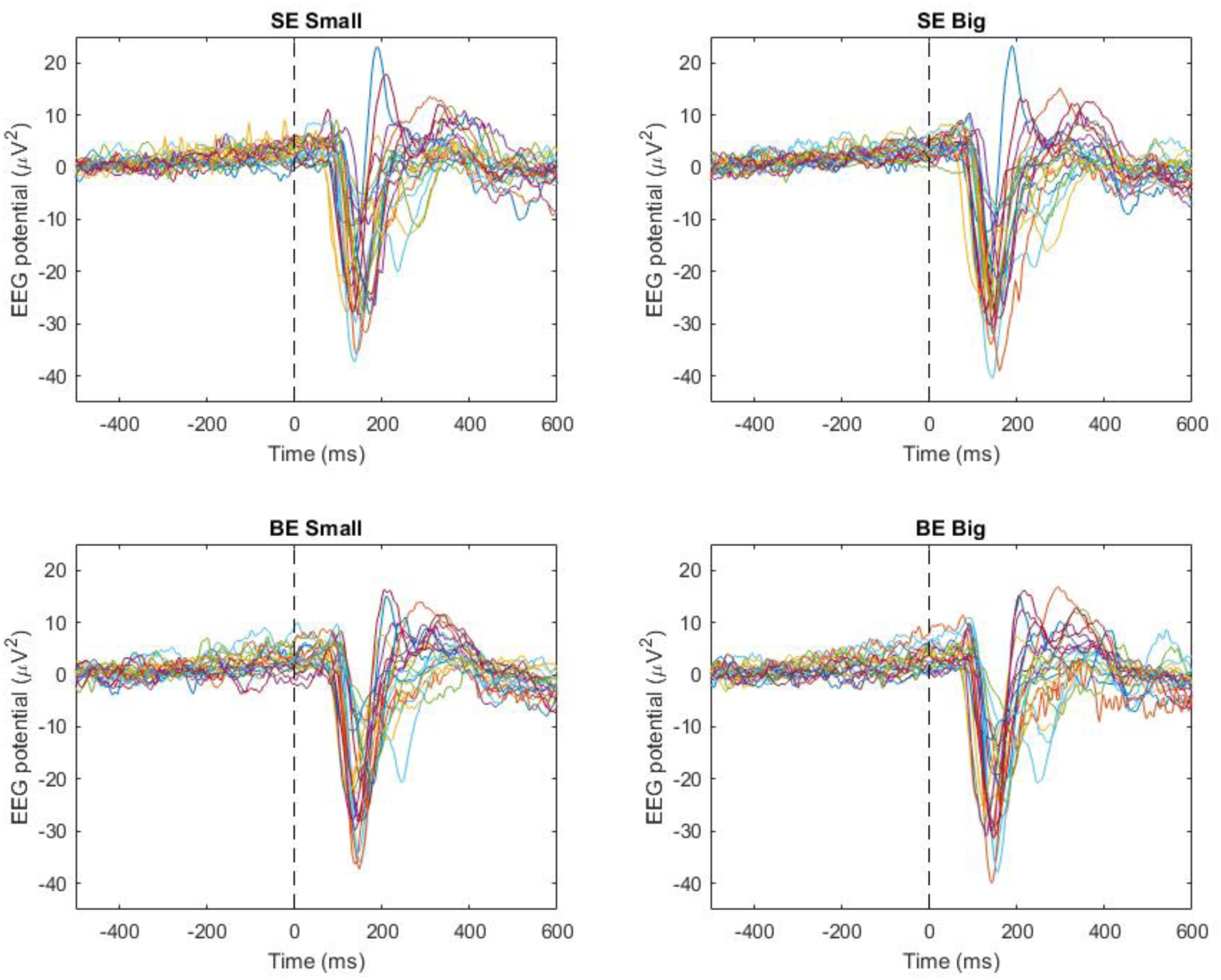
Visualisation of each participant’s grand average balance N1 waveform at Cz for each Condition x Perturbation Size combination. All ERPs are overlaid to display the between participant variability in ERP amplitude and latency following perturbation onset (vertical dashed line at time 0).

**Figure 3.3.**
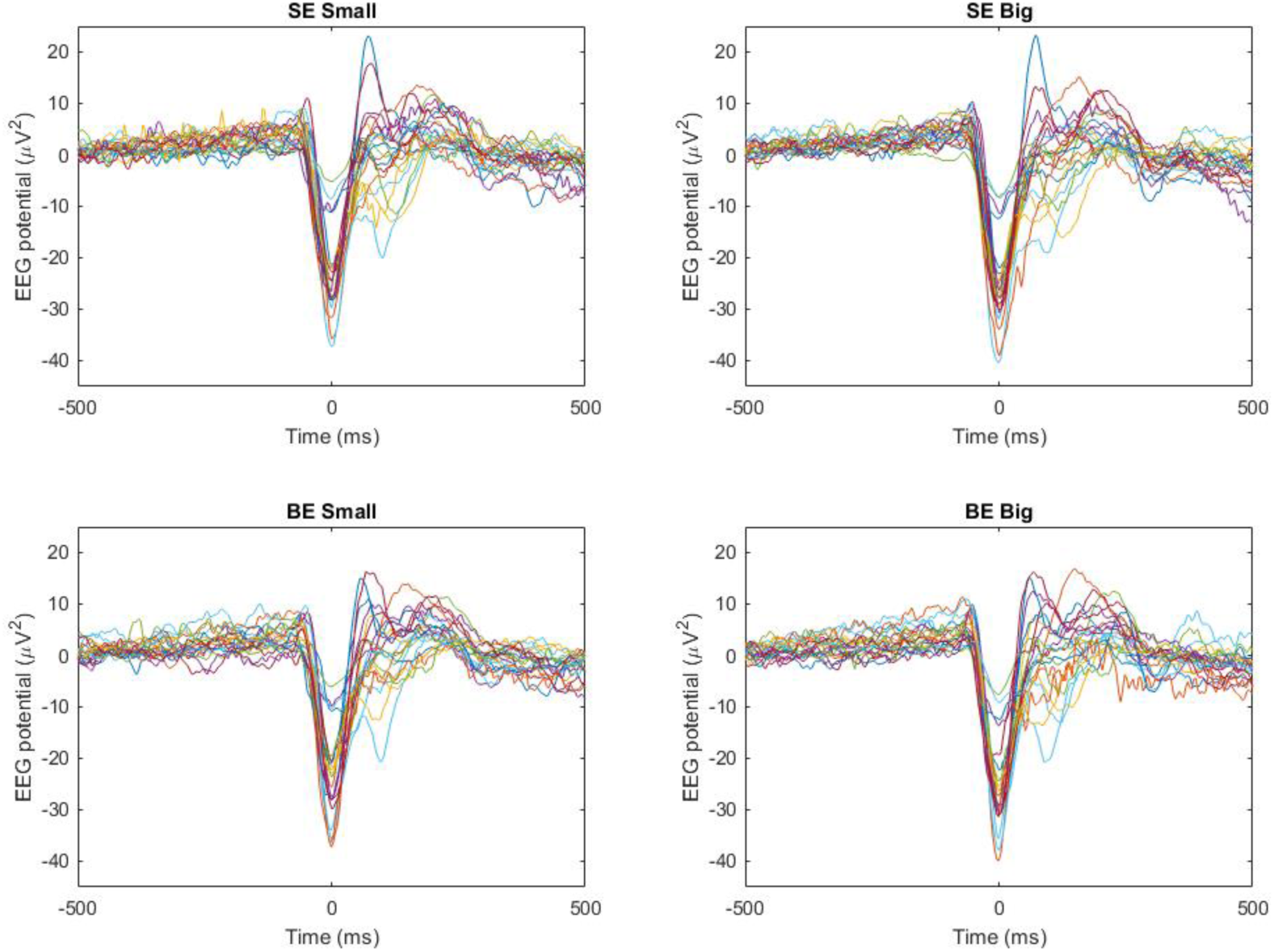
Balance N1 ERPs temporally aligned to their individual peak N1 latency (time 0). Centring each waveform on the participant-specific N1 peak reduces the influence of between-participant latency variability and allows clearer visualisation of the consistency of N1 waveform morphology across individuals. This approach highlights common features of the N1 response that may be obscured when ERPs are aligned solely to perturbation onset.

**Figure 3.4.**
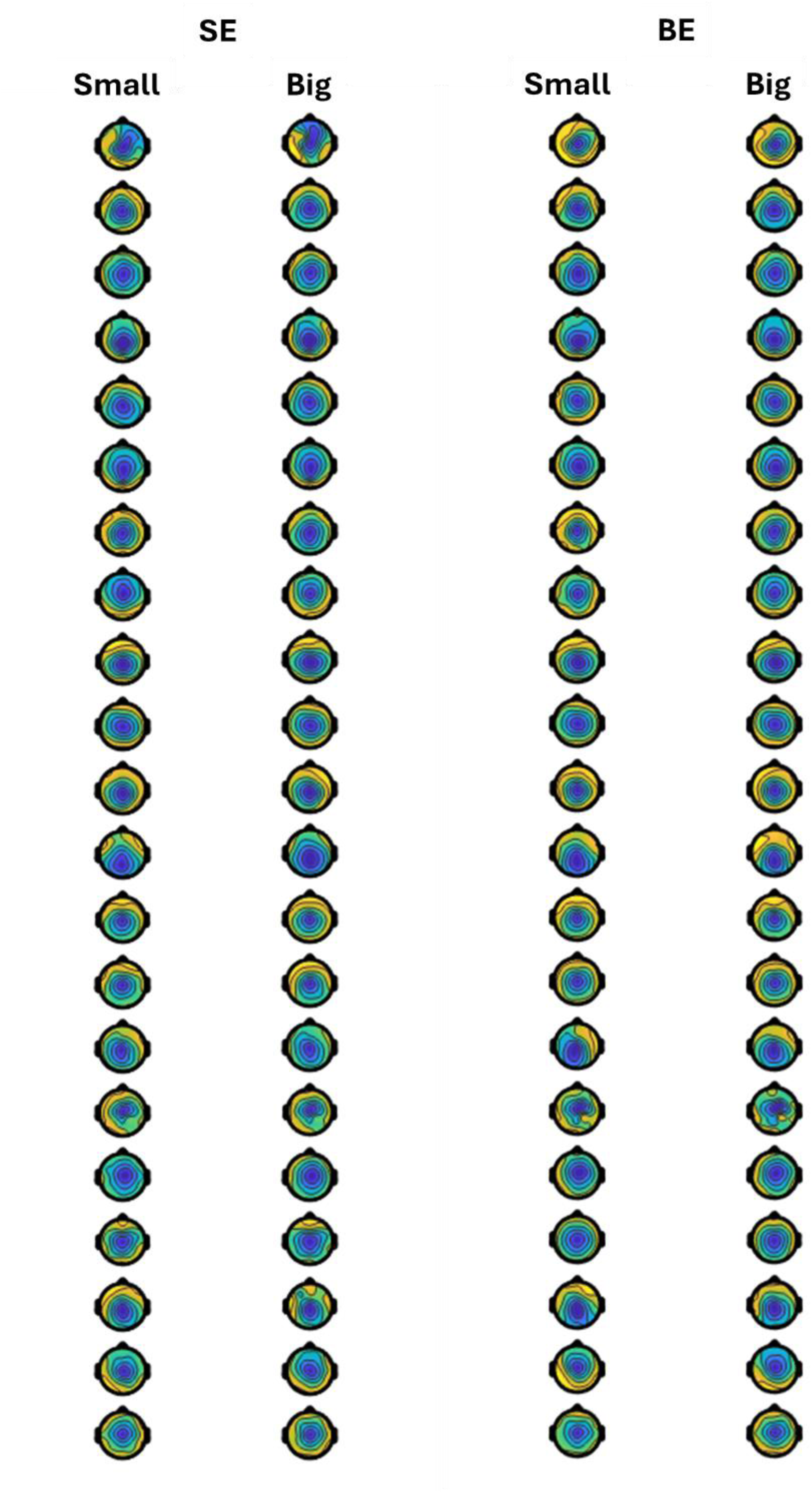
Topographical scalp maps of each participant’s average EEG activity between 100–200 ms following perturbation onset for each Condition × Perturbation Size combination. To emphasise the spatial morphology of neural activity, each participant’s scalp map is scaled independently. As a result, the maps highlight within-participant topographical patterns rather than between-participant differences in absolute amplitude.

### 4. Extended Methods

#### Selection of different trial types

During each condition (i.e., Small-Expectation and Big-Expectation), there were 45 expectation-confirming trials (e.g., small perturbations during the Small-Expectation condition) and 15 expectation-violating trials. To ensure balanced counts across trial types, we selected the 15 expectation-confirming trials for analysis (from each Expectation condition) that *immediately* preceded an expectation-violating trial (see Supplementary Figure 4 for a schematic example of the trial-selection procedure).

**Figure 4.**
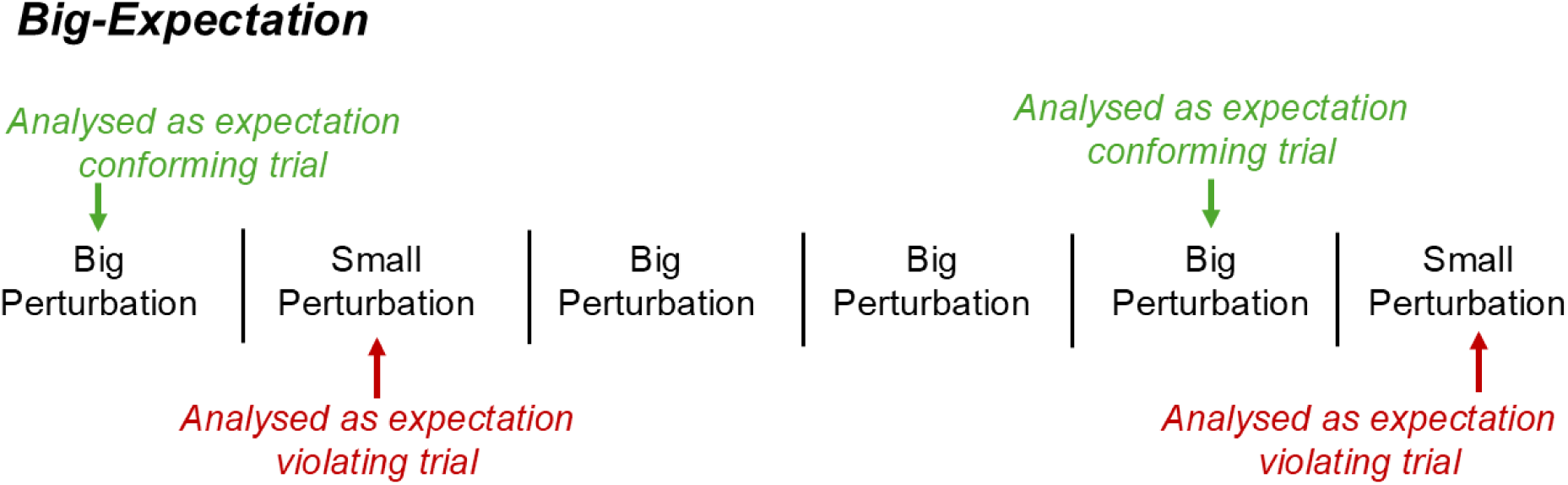
Example schematic of different perturbation trial types during the Big-Expectation condition. Note that the 15 expectation-confirming trials included in the analysis were always selected from the trial immediately preceding an expectation-violating trial.

#### EEG recording and analyses

The EEG signals were recorded at 1000 Hz from 29 active shielded AgCl electrodes embedded in a stretchable fabric cap (eego sports, ANT Neuro, Hengelo, Netherlands) positioned according to the extended 10–20 international system. Electrodes in sites CPz and AFz were used as reference and ground, respectively. Conductive gel for electrophysiological measurements was used (Signa gel, Parker), and impedance was kept below 20 kΩ. The EEG and force plate (see below) signals were synchronised through a square-wave trigger upon the initiation of an experimental recording. EEG signals were band-pass filtered using the EEGLAB “basic FIR filter (new)” (1–80 Hz, 3300 filter order, −6 dB cutoff frequency, 1 Hz transition bandwidth) prior to being cut into epochs ranging from −2 to +2 s relative to perturbation onset and re-referenced to the average of all scalp electrodes. These epochs were visually inspected for large EEG contamination from muscular artifacts, but no trials were discarded. No bad EEG channels were identified. Trials from both the Small-Expectation and Big-Expectation blocks were then stacked (i.e., 120 trials total) prior to performing independent component analysis (ICA) through the RunICA infomax algorithm(1). ICA weights that presented obvious non neural activity upon visual inspection (e.g., eyeblinks, line noise, muscular artifact) were manually rejected. On average, we retained 23.5 ± 1.9 components per participant. All processing steps were performed using EEGLAB (v2020.0) functions (2) for MATLAB.

#### Pre-perturbation EEG activity

Pre-perturbation EEG activity was analysed in a 1 s pre-perturbation window. Due to the well-established role of pre-stimulus cortical beta in modulating subsequent stimulus perception (3–5) and behavioural response (6–9), we focused our pre-perturbation EEG analyses to the beta band (15-29 Hz). Due to the transient (or, “burst-like” (5)) nature of beta activity, we chose to analyse the characteristics of individual beta burst-like events on a trial-by-trial basis, rather than averaged pre-stimulus activity. To provide a comprehensive overview of pre-perturbation beta activity, and in-line with previous research (3,4), we selected three key outcomes: beta event rate (per/s), normalised power of beta events (normalised in factors of median (FOM)), and timing of the most recent beta event (with respect to perturbation onset; ranging from -1 to 0 s). These analyses were based on the methods presented by Shin et al. (4), and were conducted using the open-source ‘SpectralEvents Toolbox’ within MATLAB (10). We used the default parameters within this toolbox (i.e., fixed-cycle complex Morlet wavelet, with 15 evenly spaced frequencies spanning 15-29 Hz). We chose a fixed-cycle width of 6 cycles (rather than the 7 cycles originally used by Shin et al. (4)) to better align this analysis with our post-perturbation time-frequency analysis; however, a replication of the analysis with 7 cycles produced identical results. Spectral events were identified as local maxima in the trial-by-trial time-frequency matrix that exceeded six times the median power of the whole time-frequency power matrix for that given electrode.

Although we hypothesised that these effects would be strongest across sensorimotor areas responsible for processing and responding to the subsequent postural perturbation (see (3–5), analyses were conducted across the whole scalp to provide a more thorough understanding of how expectation modulates the cortical processing of imbalance (with analyses adjusted for multiple comparisons; see Statistical Analyses section below).

## Funding

This research was supported by a Wellcome Trust Sir Henry Wellcome Postdoctoral Fellowship awarded to T.J.E. (Grant Number: 222747/Z/21/Z).

## Conflicts of interest

The authors declare no competing interest.

## Data availability

All data and analytical code are freely available via Open Science Framework (https://osf.io/yv2a9/).

## Notes

https://osf.io/yv2a9/

### Competing Interest Statement

The authors have declared no competing interest.

### Summary of Updates

1) Calculation and reporting of effect sizes throughout 2) New correlational analyses exploring associations between EEG outcomes and expectation-induced perceptual bias 3) Thorough expansion of the figures in the supplementary materials, including: (i) individual connected datapoints for all analyses; (ii) individual N1 traces and topographical scalp maps; (iii) figures for correlational analyses (see point #2), and; (iv) schematic outline of the experimental set-up.

