## Supplementary Materials for "Perceiving Instability: How Expectations Bias Sensorimotor Processing in Balance Control"

1. **Extended data visualisation**

The following figures are intended to illustrate the within-participant change scores that underpin the generalised estimating equation (GEE) analyses reported in the main manuscript. Each figure contains four panels showing the effect of Perturbation Size for the Small-Expectation (SE [Panels A]) and Big-Expectation (BE [Panels B]) conditions, and the effect of Expectation for both the Small (Panels C) and Big (Panels D) perturbations. Each panel contains raincloud-style plots that display individual data points connected across repeated measures with lines. Half-violin density plots and boxplots summarise data distributions, whilst diamond symbols represent the mean. The right-hand section of each panel displays participant-level difference scores (i.e., Δ = Big – Small, or Δ = BE - SE) for visualisation of within-participant effects.

**Figure S1.1.** Data visualisation of perceived instability to highlight the effect of Perturbation Size (panel A [during SE trials] and panel B [during BE trials]) and Expectation (panel C [during Small perturbations] and panel D [during Big perturbations]).

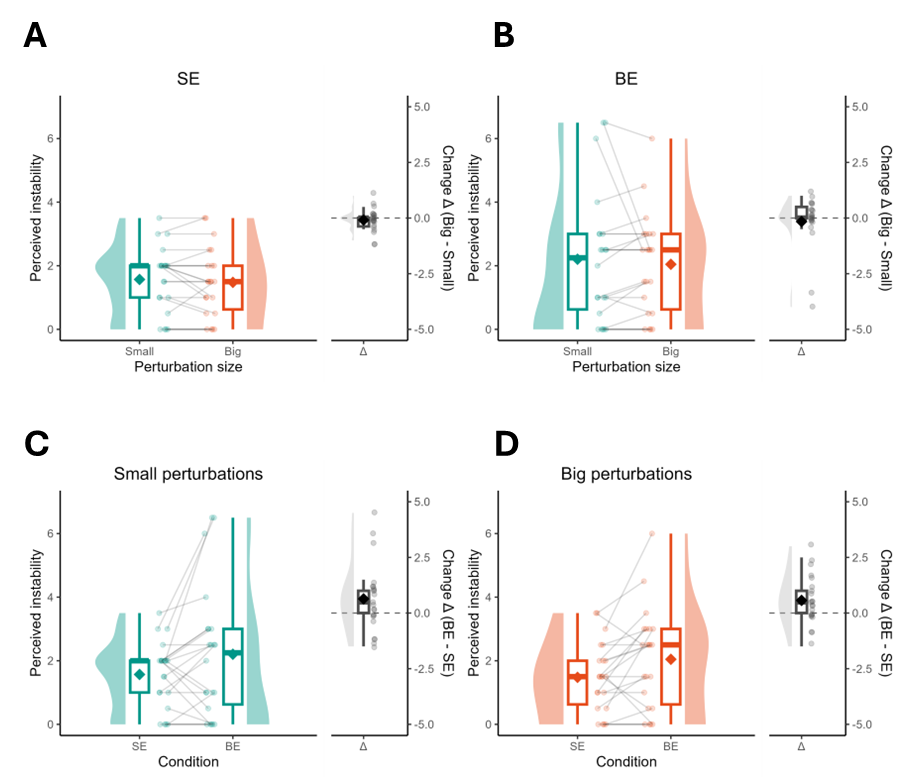

**Figure S1.2.** Data visualisation of peak backwards anteroposterior (AP) centre of pressure (CoP) velocity (m/s) to highlight the effect of Perturbation Size (panel A [during SE trials] and panel B [during BE trials]) and Expectation (panel C [during Small perturbations] and panel D [during Big perturbations]).

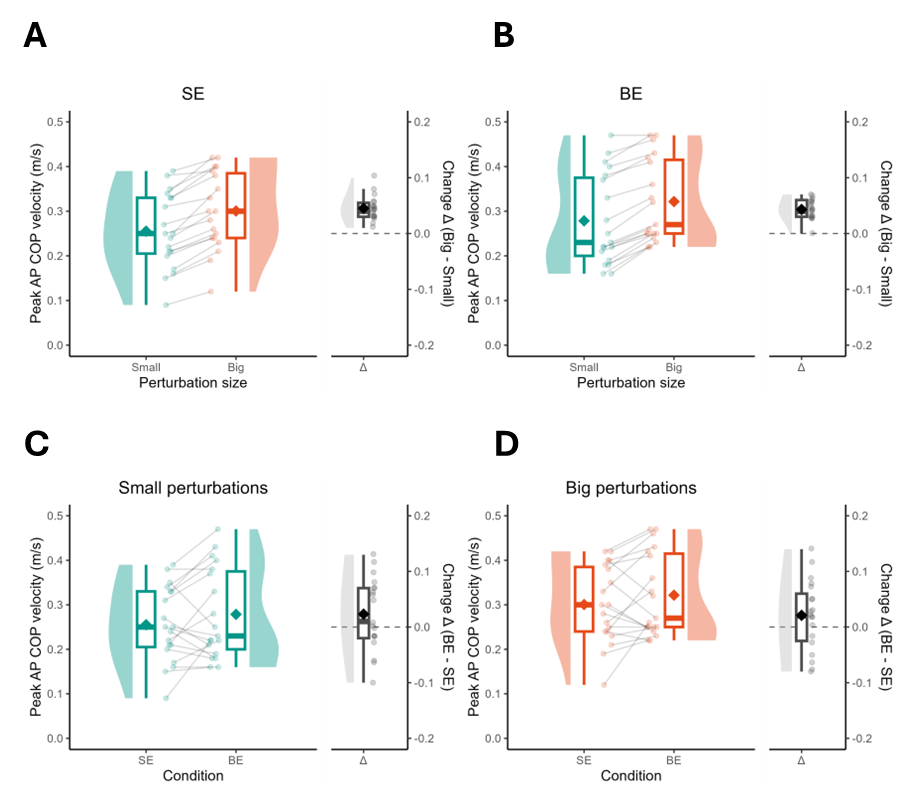

**Figure S1.3.** Data visualisation of cortical N1 amplitudes (µV) to highlight the effect of Perturbation Size (panel A [during SE trials] and panel B [during BE trials]) and Expectation (panel C [during Small perturbations] and panel D [during Big perturbations]).

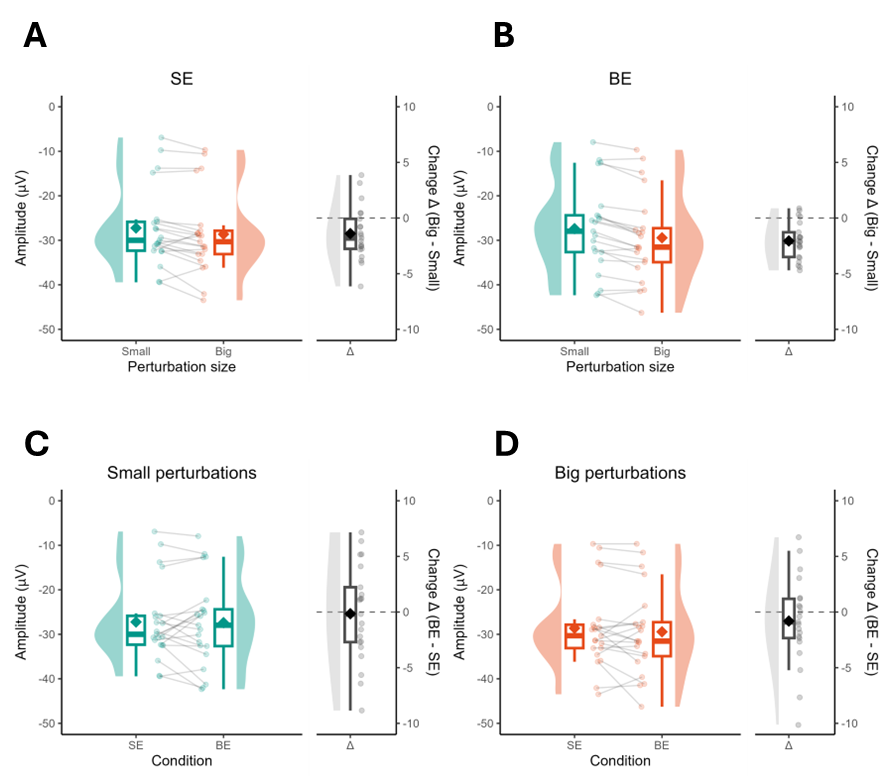

**Figure S1.4.** Data visualisation of post-perturbation (100-200 ms) beta event-related spectral power (ERSP; dB) to highlight the effect of Perturbation Size (panel A [during SE trials] and panel B [during BE trials]) and Expectation (panel C [during Small perturbations] and panel D [during Big perturbations]).

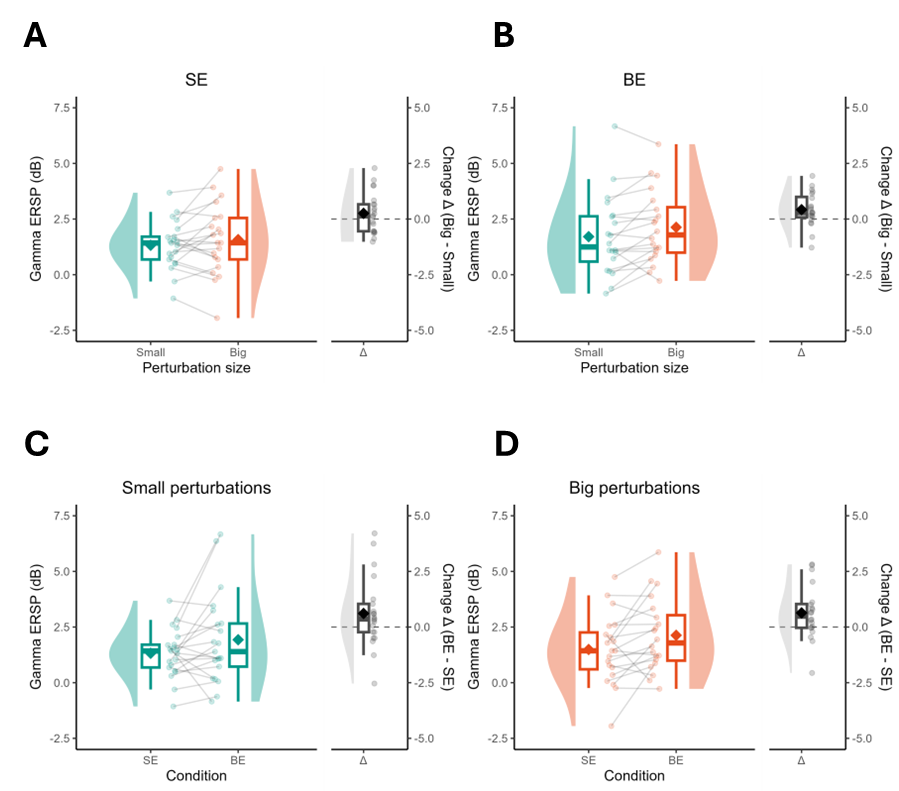

2. **Additional analyses**

As illustrated in Supplementary Figure S2.1, the timing of the final beta event (prior to perturbation onset) tended to also be earlier (i.e., further from perturbation onset) during Big-Expectation trials—specifically for frontal (F4 (*t* = 2.19, *p* = 0.040, *d* = 0.48) and FC5 (*t* = 2.79, *p* = 0.011, *d* = 0.61)) and central electrodes (C4; *t* = 2.20, *p* = 0.040, *d* = 0.48). However, these differences did not survive statistical correction for multiple comparisons.

**Figure S2.1. Topographical representation of timing of the final beta event, prior to perturbation onset**. Note, time zero indicates the time at which the perturbation occurred (i.e., an event occurring at -0.3s thus occurs 300ms prior to perturbation onset). X-markers on the t-score map indicate statistically significant differences between conditions; however, these differences did not survive statistical correction for multiple comparisons.

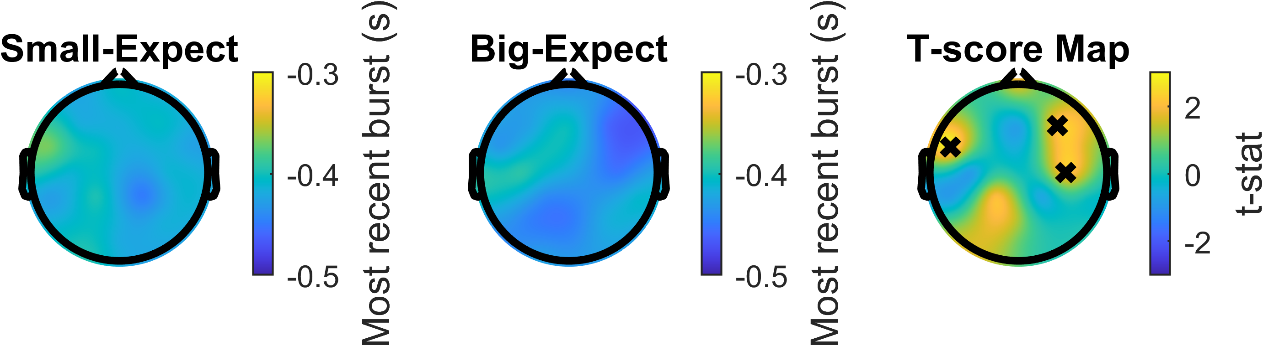

**Figure S2.2.** Scatter plot displaying the positive linear correlation between perceived instability and post-perturbation gamma ERSP (dB) during Big-Expectation (BE) trials.

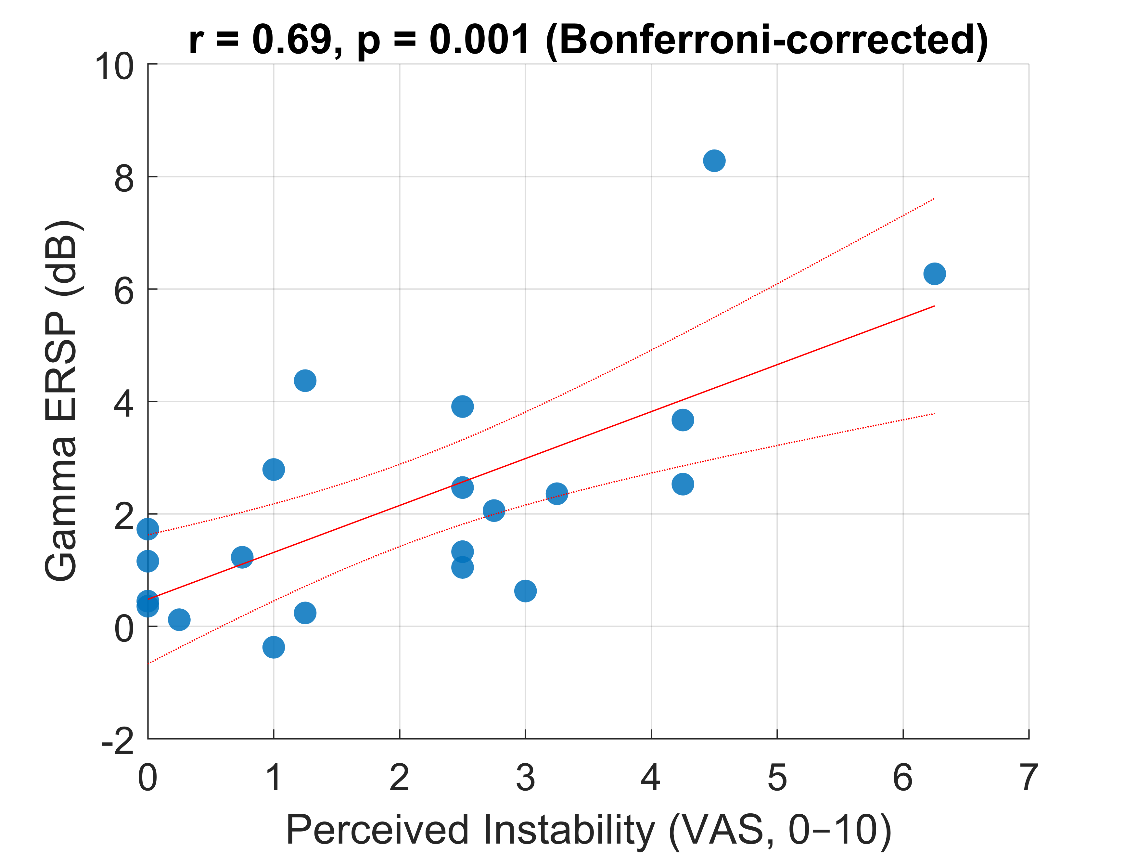

**Figure S2.3. Event-related spectral power and statistical comparisons following perturbations (exploratory analysis).** This figure shows time–frequency representations of grand average event-related spectral power (ERSP) at electrode P6 across all perturbation conditions. The first two columns depict ERSPs (in decibels) for each perturbation size (rows) and expectation condition (columns): Big perturbations are shown in the middle row, Small perturbations in the top row. Within each row, data are grouped by Expectation condition: Big-Expecting (BE; left column) and Small-Expecting (SE; middle column). Warmer colours (i.e., yellow) indicate increased spectral power relative to baseline. The third column displays pixel-wise t-value maps comparing ERSPs across Expectation conditions. For Small perturbations (top), ERSPs during the SE block are compared to those during the BE block. For Big perturbations (middle), SE-Big is compared to BE-Big. Red areas reflect greater power during BE blocks, while blue reflects greater power during SE blocks. White outlines indicate statistically significant clusters. Notably, a marked reduction in alpha activity (8–12 Hz) between 200–400 ms was observed when perturbations were larger than expected (i.e., SE-Big vs BE-Big). The adjacent topoplot shows the spatial distribution of this alpha difference, with significant effects (white circles) observed over parieto-occipital electrodes (P6 and O2), and sub-threshold effects marked with an “×” (e.g., CP5). The bottom row compares ERSPs between Big and Small perturbations within each Expectation condition (left = SE, right = BE). These contrasts revealed increased beta–gamma (25–80 Hz) activity from ~150–400 ms following larger perturbations. Corresponding topographical plots show this activity was broadly distributed across posterior electrodes.

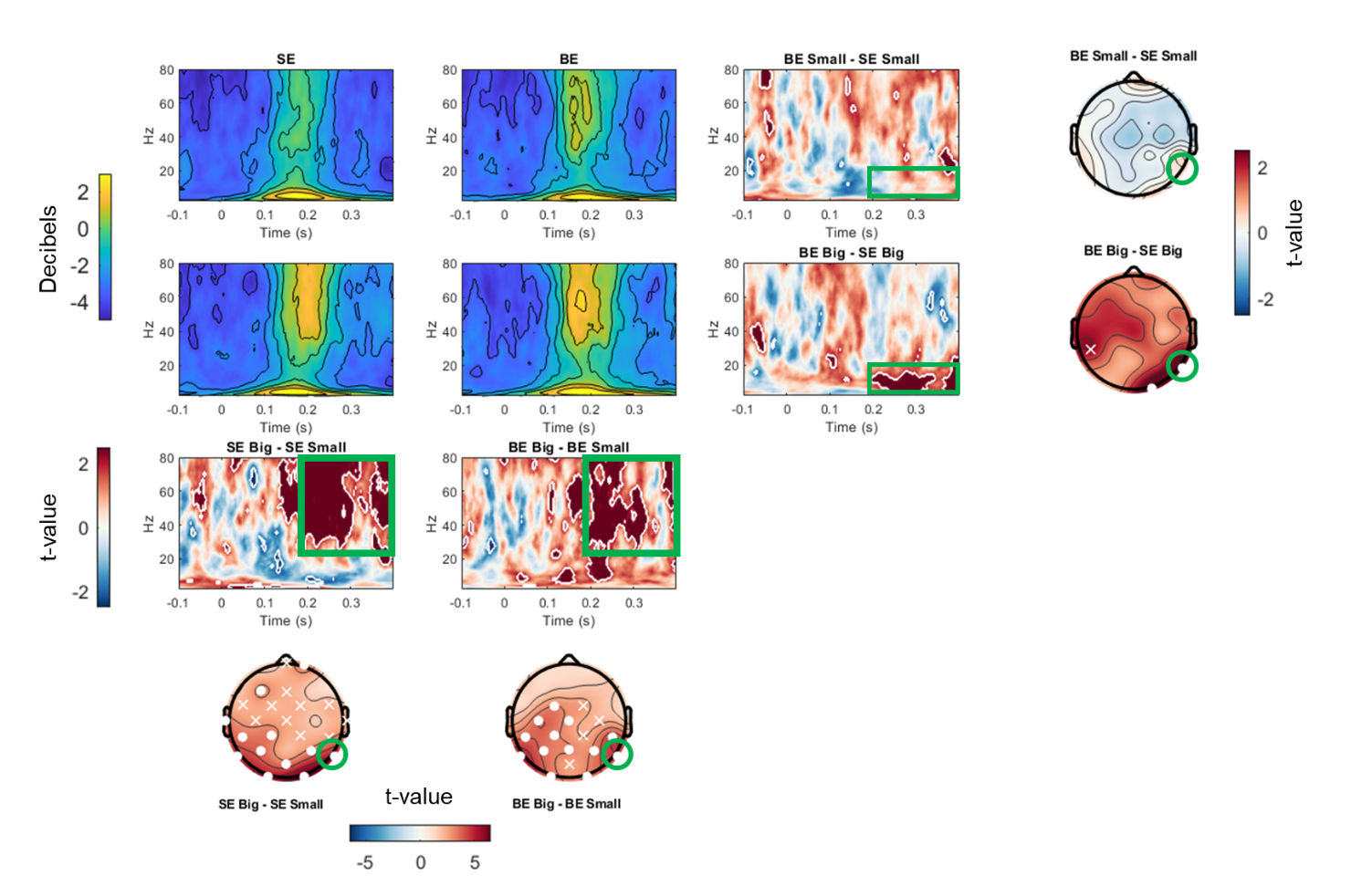

3. **Visualisation of individual balance N1 morphology**

The following supplementary figures provide complementary visualisations of the balance N1 response across Expectation and Perturbation Size. These figures are intended to illustrate individual-level variability in N1 waveform morphology, latency, and scalp topography, and to aid interpretation of the group-level analyses reported in the main manuscript. To prioritise visual clarity of within-participant patterns, several figures use participant-specific normalisation, temporal alignment, or scaling; consequently, these visualisations are descriptive and are not intended for quantitative comparison of absolute amplitudes between participants.

**Figure S3.1**. Visualisation of each participant’s grand-average balance N1 waveform at Cz for every Condition × Perturbation Size combination. The shaded yellow region marks the 50–200 ms window used to quantify peak N1 amplitude, and the vertical dashed line indicates perturbation onset (time 0). To maximise visibility of individual N1 morphology, each participant’s waveform was normalised within-participant; therefore, amplitudes are not comparable between participants.

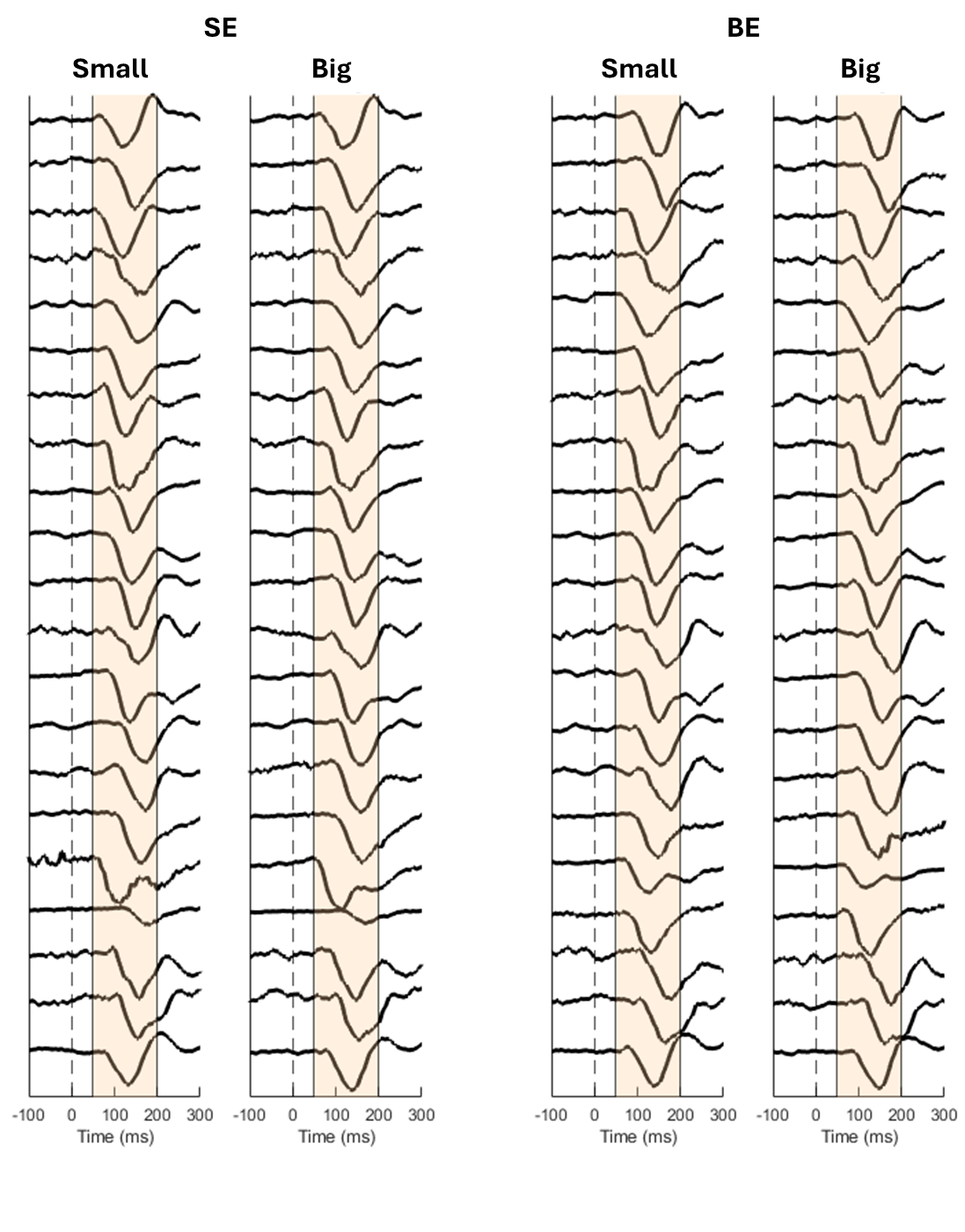

**Figure S3.2.** Visualisation of each participant’s grand average balance N1 waveform at Cz for each Condition x Perturbation Size combination. All ERPs are overlaid to display the between participant variability in ERP amplitude and latency following perturbation onset (vertical dashed line at time 0).

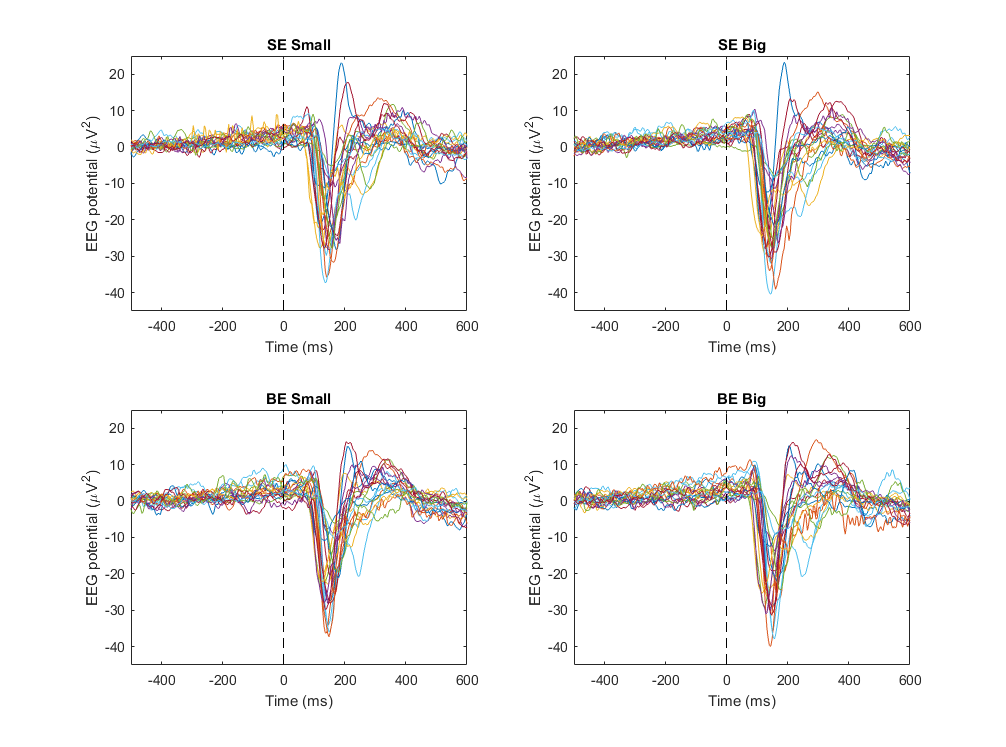

**Figure 3.3**. Balance N1 ERPs temporally aligned to their individual peak N1 latency (time 0). Centring each waveform on the participant-specific N1 peak reduces the influence of between-participant latency variability and allows clearer visualisation of the consistency of N1 waveform morphology across individuals. This approach highlights common features of the N1 response that may be obscured when ERPs are aligned solely to perturbation onset.

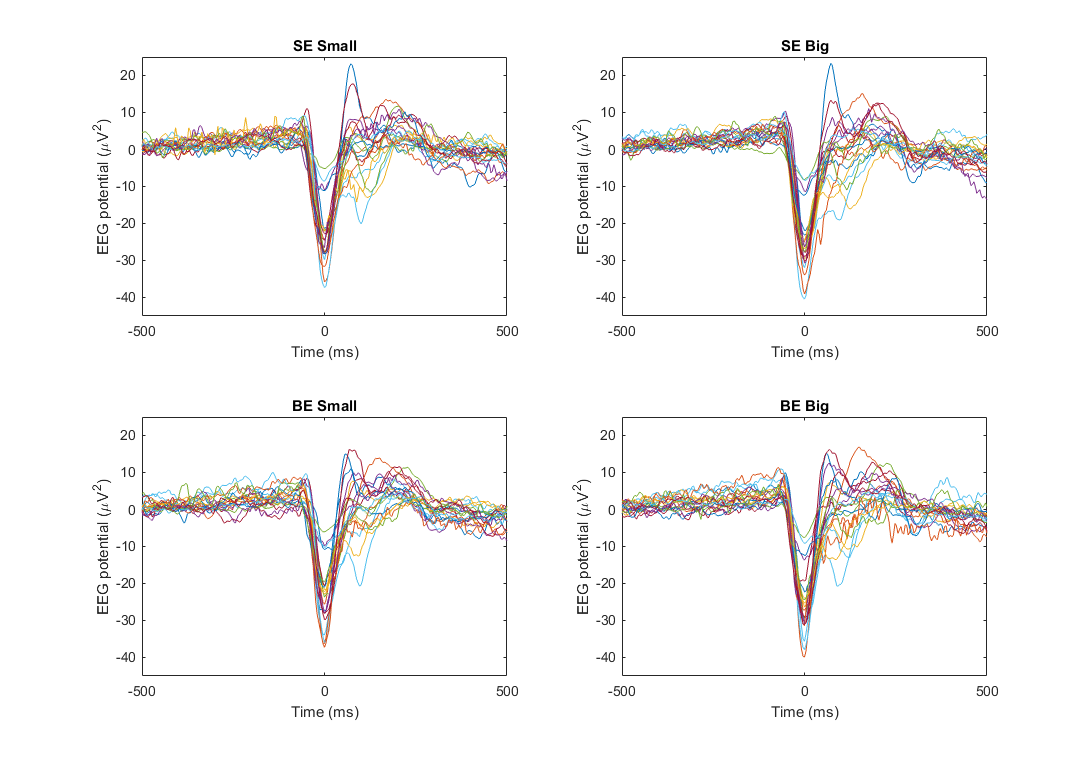

**Figure 3.4.** Topographical scalp maps of each participant’s average EEG activity between 100–200 ms following perturbation onset for each Condition × Perturbation Size combination. To emphasise the spatial morphology of neural activity, each participant’s scalp map is scaled independently. As a result, the maps highlight within-participant topographical patterns rather than between-participant differences in absolute amplitude.

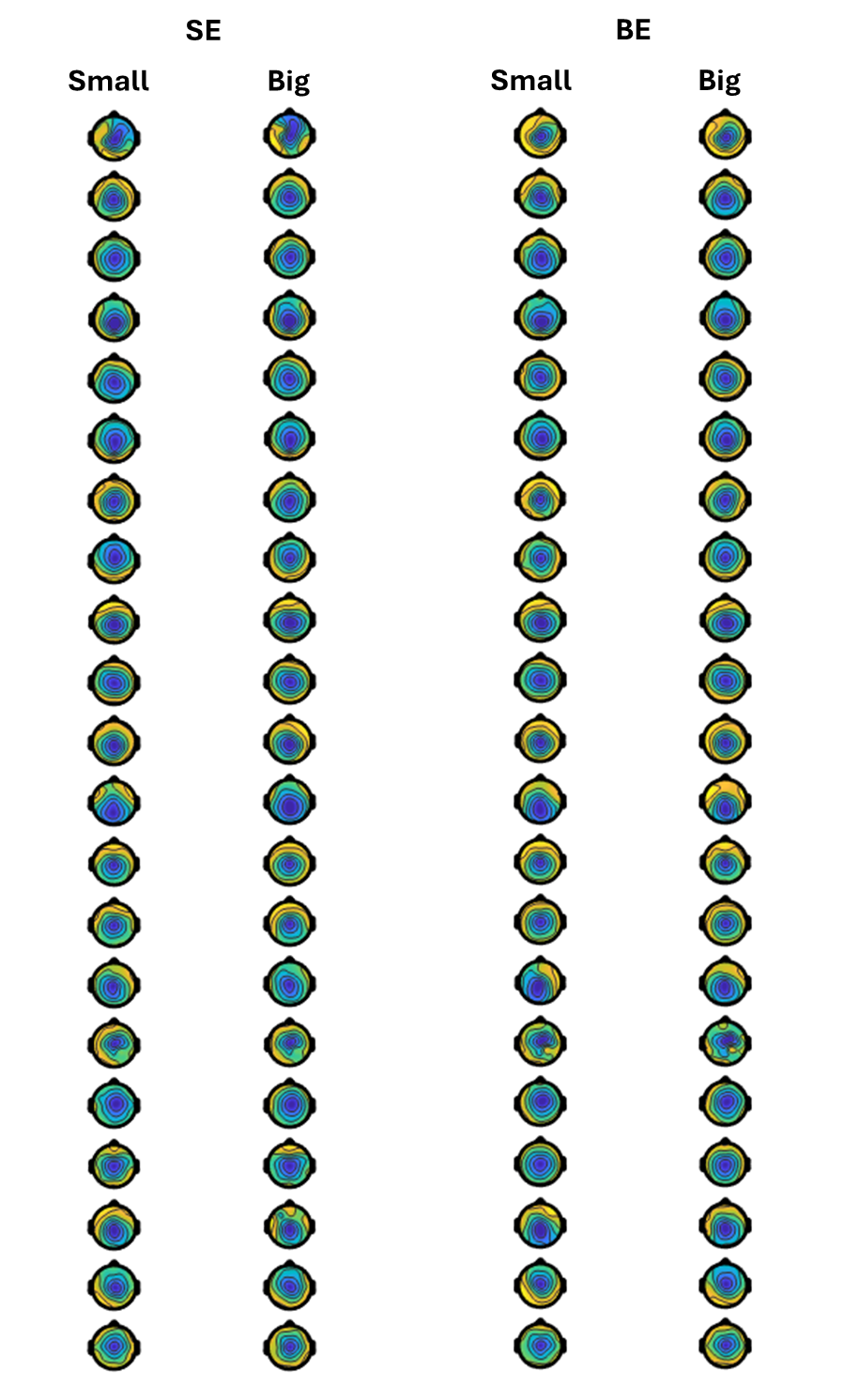

4. **Extended Methods**

**Selection of different trial types**

During each condition (i.e., Small-Expectation and Big-Expectation), there were 45 expectation-confirming trials (e.g., small perturbations during the Small-Expectation condition) and 15 expectation-violating trials. To ensure balanced counts across trial types, we selected the 15 expectation-confirming trials for analysis (from each Expectation condition) that *immediately* preceded an expectation-violating trial (see Supplementary Figure 4 for a schematic example of the trial-selection procedure).

**Figure 4.** Example schematic of different perturbation trial types during the Big-Expectation condition. Note that the 15 expectation-confirming trials included in the analysis were always selected from the trial immediately preceding an expectation-violating trial.

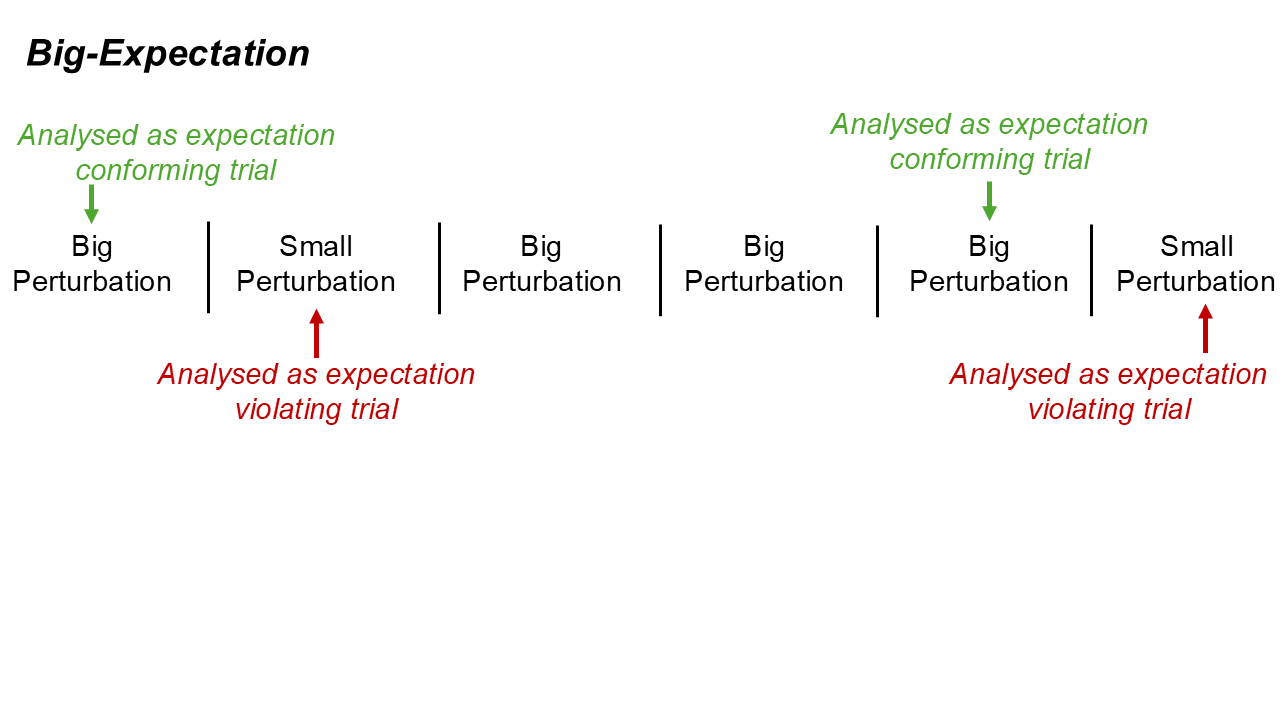

**EEG recording and analyses**

The EEG signals were recorded at 1000 Hz from 29 active shielded AgCl electrodes embedded in a stretchable fabric cap (eego sports, ANT Neuro, Hengelo, Netherlands) positioned according to the extended 10–20 international system. Electrodes in sites CPz and AFz were used as reference and ground, respectively. Conductive gel for electrophysiological measurements was used (Signa gel, Parker), and impedance was kept below 20 kΩ. The EEG and force plate (see below) signals were synchronised through a square-wave trigger upon the initiation of an experimental recording. EEG signals were band-pass filtered using the EEGLAB “basic FIR filter (new)” (1–80 Hz, 3300 filter order, −6 dB cutoff frequency, 1 Hz transition bandwidth) prior to being cut into epochs ranging from −2 to +2 s relative to perturbation onset and re-referenced to the average of all scalp electrodes. These epochs were visually inspected for large EEG contamination from muscular artifacts, but no trials were discarded. No bad EEG channels were identified. Trials from both the Small-Expectation and Big-Expectation blocks were then stacked (i.e., 120 trials total) prior to performing independent component analysis (ICA) through the RunICA infomax algorithm(1). ICA weights that presented obvious non neural activity upon visual inspection (e.g., eyeblinks, line noise, muscular artifact) were manually rejected. On average, we retained 23.5 ± 1.9 components per participant. All processing steps were performed using EEGLAB (v2020.0) functions (2) for MATLAB.

5. **References**

10.	jonescompneurolab/SpectralEvents. Available from: https://github.com/jonescompneurolab/SpectralEvents
